# YAP and TAZ promote periosteal osteoblast precursor expansion and differentiation for fracture repair

**DOI:** 10.1101/2020.03.17.995761

**Authors:** Christopher D. Kegelman, Madhura P. Nijsure, Yasaman Moharrer, Hope B. Pearson, James H. Dawahare, Kelsey M. Jordan, Ling Qin, Joel D. Boerckel

**Author notes:** To whom correspondence should be addressed, Mailing Address: Joel D. Boerckel, 376A Stemmler Hall, 3450 Hamilton Walk, University of Pennsylvania, Philadelphia, PA, 19104-6081.

## Abstract

In response to bone fracture, periosteal progenitor cells proliferate, expand, and differentiate to form cartilage and bone in the fracture callus. These cellular functions require the coordinated activation of multiple transcriptional programs, and the transcriptional regulators Yes-associated protein (YAP) and transcriptional co-activator with PDZ-binding motif (TAZ) regulate osteochondroprogenitor activation during endochondral bone development. However, recent observations raise important distinctions between the signaling mechanisms used to control bone morphogenesis and repair. Here, we tested the hypothesis that YAP and TAZ regulate osteochondroprogenitor activation during endochondral bone fracture healing. Constitutive YAP and/or TAZ deletion from Osterix-expressing cells impaired both cartilage callus formation and subsequent mineralization. However, this could be explained either by direct defects in osteochondroprogenitor differentiation after fracture, or by developmental deficiencies in the progenitor cell pool prior to fracture. Consistent with the second possibility, we found that developmental YAP/TAZ deletion produced long bones with impaired periosteal thickness and cellularity. Therefore, to remove the contributions of developmental history, we next generated adult onset-inducible knockout mice (using Osx1-Cre^tetOff^) in which YAP and TAZ were deleted prior to fracture, but after normal development. Adult onset-induced YAP/TAZ deletion had no effect on cartilaginous callus formation, but impaired bone formation at 14 days post-fracture (dpf). Earlier, at 4 dpf, adult onset-induced YAP/TAZ deletion impaired the proliferation and expansion of osteoblast precursor cells located in the shoulder of the callus. Further, activated periosteal cells isolated from this region at 4 dpf exhibited impaired osteogenic differentiation *in vitro* upon YAP/TAZ deletion. Finally, confirming the effects on osteoblast function *in vivo*, adult onset-induced YAP/TAZ deletion impaired bone formation in the callus shoulder at 7 dpf, prior to the initiation of endochondral ossification. Together, these data show that YAP and TAZ promote the expansion and differentiation of periosteal osteoblast precursors to accelerate bone fracture healing.

## INTRODUCTION

Bone is a remarkable tissue in its capacity to heal without forming a scar, and most bone injuries heal readily, with bone fractures healing at success rates of 90-95%^(1)^. This is due, in part, to the maintenance of a skeletal stem cell population capable of recapitulating many aspects of embryological programs for adult tissue regeneration^(2)^. As in development, bone formation during fracture repair can occur through both direct intramembranous ossification and endochondral ossification via a cartilage intermediate^(3–5)^. However, the source, niche, and molecular regulation of the progenitor cells responsible for bone fracture repair are distinct from those that produce the skeleton during development^(2,6)^. In the embryo, mesodermal mesenchymal progenitors in the limb bud form a template of the skeletal elements, while bone fracture healing initiates by expansion and differentiation of osteochondroprogenitor cells resident in the bone-lining periosteum^(3,4,7)^. Understanding the mechanisms that regulate the proliferation and differentiation of these cells will be critical to develop new therapeutic strategies for accelerating fracture repair and regenerating challenging bone injuries that cannot heal on their own.

To define the roles of the molecular mediators that regulate adult periosteal progenitor cell function, we must decouple the developmental history of the osteochondroprogenitor cells that come to reside in the periosteum from the regenerative function of these cells after fracture to accurately evaluate their contribution to responding to injury in the adult. For example, a recent study in which PDGFRβ was deleted from Osterix-expressing cells found no notable defects in skeletal development, but severe impairment of fracture healing^(8)^, demonstrating contextual distinctions between development and fracture repair. Further, skeletal cell- and bone-specific gene deletion during development may alter the number, location, or niche of progenitor cells that, during injury, are activated for skeletal regeneration^(9)^. In this study, we assessed the effects of conditional gene deletion from osteochondroprogenitors on endochondral bone fracture repair, with deletion performed either constitutively during development or inducibly after normal development to skeletal maturity prior to fracture. Several types of inducible Cre-loxP systems exist to temporally regulate Cre-mediated gene recombination, including the interferon-responsive *Mx1* promoter^(10)^, mifepristone^(11)^, tamoxifen-inducible mutated estrogen- and progesterone-receptors^(12)^, and tetracycline-controlled systems^(13)^. Here, we used the Osterix-Cre^tetOff^ (Osx-Cre) mouse in which Cre-recombinase is driven by the Osterix promoter and temporally controlled by tetracycline (or its more stable derivative, doxycycline)^(14)^. Osx-Cre is expressed in hypertrophic chondrocytes and osteoprogenitors, including those of the primary ossification center in the embryo as well as the periosteum in the adult^(14,15)^.

Periosteal cell expansion and differentiation require the coordinated activation of multiple transcriptional programs, and the transcriptional regulators, Yes-associated protein (YAP) and transcriptional co-activator with PDZ-binding motif (TAZ) have recently emerged as critical mediators of osteoblast progenitor proliferation and differentiation during bone development^(16–18)^. Previously, we found that constitutive homozygous deletion of both YAP and TAZ from Osterix-expressing cells caused severe skeletal fragility and neonatal lethality^(16)^. Mice with a single allele of either YAP or TAZ survived, but sustained spontaneous long bone fractures due to both reduced bone mass and defects in bone matrix collagen that caused weaker bone mechanical properties^(16)^. Despite fracture, the neonatal limbs exhibited natural reduction^(19)^ and eventually healed through callus formation ^(16)^. However, the roles of YAP and TAZ in periosteal progenitor cell function and their contributions to bone fracture healing are unknown.

Here, we conditionally deleted YAP and/or TAZ from Osterix-expressing cells using either constitutive or tetOff-inducible deletion and evaluated adult endochondral bone fracture healing. We found that constitutive YAP and/or TAZ deletion impaired both callus formation and subsequent mineralization, due to developmental deficiencies in the progenitor cell pool prior to fracture. In contrast, adult onset-induced YAP/TAZ deletion had no effect on cartilaginous callus formation, but impaired both the proliferation of osteoblast precursor cells located in the shoulder of the callus and their osteogenic differentiation, both, *in vitro* and *in vivo*. Together, these data show that YAP and TAZ promote the expansion and differentiation of periosteal osteoblast precursors to accelerate bone fracture healing.

## MATERIALS AND METHODS

### Animals

Mice harboring loxP-flanked exon 3 alleles in both YAP and TAZ on a mixed C57BL/6J genetic background were kindly provided by Dr. Eric Olson (University of Texas Southwestern Medical Center). Tetracycline responsive B6.Cg-Tg(Sp/7-tTA,tetO-EGFP/Cre)1AMc/J (Osx-Cre^tetOff^) mice from The Jackson Laboratory (Bar Harbor, MA, USA) were used to generate two mouse models in which we conditionally deleted YAP and/or TAZ from Osterix-expressing cells (**Table 1**). In both mouse models, tetracycline (or its more stable derivative, doxycycline) administration prevents tetracycline-controlled transactivator protein (tTA) binding to the tetracycline-responsive promoter element (TRE) in the promoter of the Cre transgene, allowing Cre expression only in the absence of doxycycline^(14)^ for temporal control of Osx-Cre-mediated gene deletion.

**Table 1:** Experimental fracture healing models, genotypes, and abbreviations.

| Deletion approach | Genotype | Abbreviation |
| --- | --- | --- |
| Constitutive | Yap<sup>fl/fl</sup>;Taz<sup>fl/fl</sup> | YAP<sup>WT</sup>;TAZ<sup>WT</sup> |
|  | Yap<sup>fl/+</sup>;Taz<sup>fl/+</sup>;Osx-Cre | YAP<sup>cHET</sup>;TAZ<sup>cHET</sup> |
|  | Yap<sup>fl/fl</sup>;Taz<sup>fl/+</sup>;Osx-Cre | YAP<sup>cKO</sup>;TAZ<sup>cHET</sup> |
|  | Yap<sup>fl/+</sup>;Taz<sup>fl/fl</sup>;Osx-Cre | YAP<sup>cHET</sup>;TAZ<sup>cKO</sup> |
|  | Yap<sup>fl/fl</sup>;Taz<sup>fl/fl</sup>;Osx-Cre | YAP<sup>cKO</sup>;TAZ<sup>cKO</sup> |
| Inducible | Osx-Cre<sup>tetOff</sup> | Osx:Cre |
|  | Yap<sup>fl/fl</sup>;Taz<sup>fl/fl</sup> | YAP<sup>WT</sup>;TAZ<sup>WT</sup> |
|  | Yap<sup>fl/fl</sup>;Taz<sup>fl/fl</sup>;Osx-Cre<sup>tetOff</sup> | YAP<sup>cKOi</sup>;TAZ<sup>cKOi</sup> |

In our first study, we evaluated constitutive allele dose-dependent deletion of YAP and/or TAZ in skeletally mature mice 16-21 weeks of age (**Table 1**). Mice with homozygous floxed alleles for both YAP and TAZ (YAP^fl/fl^;TAZ^fl/fl^) were mated with double heterozygous conditional knockout mice (YAP^fl/+^;TAZ^fl/+^;Osx-Cre) to produce eight possible genotypes in each litter, but only Cre-positive and YAP^fl/fl^;TAZ^fl/fl^ animals were compared (**Table 1**). Here, the littermate YAP^fl/fl^;TAZ^fl/fl^ mice were considered the wild-type control genotype. All of these mice were bred, raised, and evaluated without tetracycline administration to induce gene recombination in Osx1-Cre-expressing cells prior to embryonic development, for the duration of the analyses in the constitutive deletion model.

In our second study, we allowed mice to develop to skeletal maturity and induced homozygous YAP/TAZ deletion two weeks prior to fracture at 16-18 weeks of age (**Table 1**). In the inducible deletion model, both littermate YAP^fl/fl^;TAZ^fl/fl^ (YAP^WT^;TAZ^WT^) mice and separately-bred Osx-Cre^tetOff^ mice were evaluated as wild type control genotypes (**Table 1**). All mice were bred and raised until skeletal maturity with doxycycline in their drinking water to prevent Cre-mediated gene recombination. For all *in vivo* fracture healing assessments, doxycycline was removed two weeks prior to fracture surgery and normal drinking water was provided for the remainder of the study. For periosteal progenitor cell isolations from fractured limbs, doxycycline was provided for both YAP^WT^;TAZ^WT^and YAP^cKOi^;TAZ^cKOi^ mice throughout skeletal development and the duration of the fracture healing experiment.

In both studies, mice were tail or ear clipped after weaning or prior to euthanasia and genotyped by an external service (Transnetyx, Inc.). All mice were fed regular chow (PicoLab Rodent Diet, Cat#: 0007688, LabDiet) *ad libitum* and housed in cages containing 2-5 animals each. Mice were maintained at constant 25°C on a 12-hour light/dark cycle. Both male and female mice were evaluated with the same fracture healing procedure for both the constitutive and inducible deletion models of fracture healing **(Table 1**). All protocols were approved by the Institutional Animal Care and Use Committees at the University of Notre Dame and the University of Pennsylvania. All animal procedures were performed in adherence to federal guidelines for animal care and conform to the Animal Research: Reporting of In Vivo Experiments (ARRIVE) guidelines.

### Open femoral fracture model and timepoints

An open, unilateral, and intramedullary pin-stabilized femoral fracture model was used to study bone repair in both the inducible and constitutive deletion studies. Femora were surgically exposed and manually fractured by applying a bending moment^(20)^, but stabilized with an intramedullary pin^(21)^. For the surgical procedure, animals were anesthetized using isoflurane (1-5%), all hair was removed from the surgical site, and the area was cleansed with sterile water followed by betadine. A 25-gauge needle was inserted in a retrograde manner into the intramedullary canal of the right femur^(22)^. Subsequently, the muscle surrounding the same femur was blunt dissected to expose the femoral midshaft and a reproducible fracture was created by applying a three-point bending moment in the femur containing the intramedullary pin. The contralateral leg was left intact. Any animals that displayed intramedullary pin displacement or fractures that were comminuted or too oblique were removed. The mice were allowed to recover under a heating lamp and after awakening returned to their cages and allowed to ambulate freely. In the constitutive deletion model, mice were euthanized at 14 days post-fracture (dpf) and 42 dpf. In the inducible deletion model, mice were euthanized at 4 dpf, 7 dpf, and 14 dpf. At 4 dpf, mice were injected intraperitoneally with 5-ethynyl-2’-deoxyuridine (EdU; E10187; Invitrogen) at 10 mg/kg 3 hours prior to euthanasia to assay cellular proliferation.

### Microcomputed tomography

Micro-computed tomography (microCT) was performed according to published guidelines^(23)^ on two separate systems. In the constitutive deletion study, 14 and 42 dpf fractured limbs were dissected free from surrounding musculature and the intramedullary pins were removed. Samples from 14 and 42 dpf limbs were wrapped in PBS-soaked gauze and frozen at −20°C. When removed from the freezer, bones were allowed to thaw while being imaged using a μCT 35 system (Scanco Medical). Samples from 14 and 42 dpf bone were imaged with an X-ray intensity of 114 μA, energy of 70 kVp, integration time of 200 ms, and resolution of 15μm. Based on the precedent set in a similar fracture healing study^(22,24)^, we defined the fracture callus mineralization threshold as 50% of the mineral density that we used to segment intact cortical bone under these conditions on this system. 2D tomograms of the fracture calluses both excluding and including the intact cortical bone were manually contoured, stacked and binarized by applying a Gaussian filter (sigma =0.8, support =1) at a threshold of 345 mg HA/cm^3^. 3D quantification of the mineralized callus excluding the intact bone were reported as total callus volume, percent callus mineralization, and volumetric bone mineral density. Five to eight mice were analyzed per group. Investigators were blinded to animal genotype during scan quantification.

In the inducible deletion study, 7 and 14 dpf limbs were dissected free from surrounding musculature and the intramedullary pins were removed. Bones from 7 and 14 dpf were then snap-frozen in liquid nitrogen-cooled isopentane for 1-minute, wrapped in gauze and imaged on a vivaCT 80 system (Scanco Medical). Samples from 7 and 14 dpf were imaged with an X-ray intensity of 114 μA, energy of 70 kVp, integration time of 200 ms, and resolution of 15μm. We again defined the fracture callus mineralization threshold as 50% of the mineral density that we used to segment intact cortical bone under these conditions on this system. 2D tomograms of the fracture calluses both excluding and including the intact cortical bone were manually contoured, stacked and binarized by applying a Gaussian filter (sigma =0.8, support =1) at a threshold of 254 mg HA/cm^3^. 3D quantification of the mineralized callus excluding the intact bone were reported as total callus volume, percent callus mineralization, and volumetric bone mineral density. Six to eight mice were analyzed per group. Investigators were blinded to animal genotype during scan quantification.

To assess the cross-sectional bone distributions on the scans from both systems, the “Bone Midshaft” evaluation script (Scanco Medical) was used to quantify polar moment of inertia (pMOI)^(25)^. Limbs from 14 dpf in the constitutive deletion model and 14 dpf from the inducible deletion model were evaluated on each system using a negative, non-physiological threshold so as to include all parts of the callus in the “Bone Midshaft” evaluation. Limbs from 42 dpf were scanned on the μCT 35 system were evaluated using the same threshold for mineralized tissue above (345 mg HA/cm^3^). In both cases, pMOI values for all groups were binned into 25 equal distance bins from the center of the fracture using a custom MATLAB script and presented as the mean with error bars corresponding to the standard deviation (SD).

### Mechanical testing

Following microCT scanning, 14 and 42 dpf limbs from the constitutive deletion model were tested in torsion to failure. For torsional testing, we used fixtures and a custom potting apparatus that allowed us to reproducibly align and pot the fractured limbs in polymethylmethacrylate bone cement. After the fractured limbs were potted, they were loaded in torsion at a rate of 1°/s until failure using a custom-designed micro-torsional testing system. Recorded torque-rotation data were normalized by gauge length on a per-sample basis^(26)^. Torsional rigidity, maximum torque to failure, work to maximum torque and work to failure were quantified from the normalized torque-rotation data using a custom MATLAB script^(16)^. Five to eight mice were analyzed per group per timepoint. Investigators were blinded to animal genotype during data quantification.

### Histology, immunohistochemistry, and immunofluorescence

Limbs from 7 and 14 dpf were fixed with 10% neutral buffered formalin for 48 hours and decalcified for 4 weeks with 0.25M EDTA (pH 7.4) at 4°C. Paraffin sections (5 μm thickness) were processed for either immunohistochemistry or histology. Primary antibodies were compared to normal rabbit sera IgG control sections. For immunostaining, anti-OSX (1:500, ab22552; abcam), anti-YAP (1:500, 14074; Cell Signaling), and anti-TAZ (1:250 NB110-58359; Novus Biologicals) primary antibodies were applied overnight. Next, sections were incubated with corresponding biotinylated secondary antibody, avidin-conjugated peroxidase, and diaminobenzidine substrate chromogen system (329ANK-60; Innovex Biosciences), which allowed for immunohistochemical detection of positively stained cells. Hematoxylin and eosin stains (H&E), Safranin-O, and Picrosirius Red stains were used to stain for bone, cartilage, and collagen.

Limbs from 4 dpf and intact femora were fixed with 10% neutral buffered formalin for 48 hours at 4°C, transferred to 30% sucrose in PBS overnight at 4°C, and then embedded in O.C.T. compound (Tissue-Tek). Thin sections (7 μm thickness) were made from undecalcified fractured femurs using cryofilm IIC tape (Section Lab Co. Ltd.) as previously described^(27)^ and processed for immunofluorescence and/or aqueous H&E staining. Taped sections were glued to microscope slides using a UV-adhesive glue, rehydrated and then decalcified with 0.25M EDTA (pH 7.4) for 3 minutes prior to staining. 5-ethynyl-2’-deoxyuridine (EdU)) staining were performed using the Click-iT Plus EdU Assay kit (C10339; Invitrogen) according to the manufacturer’s instructions.

### Imaging and histomorphometric analysis

Histological and immunohistochemical sections were imaged on either on an Axio Observer Z1 (Zeiss) at the 10x and 25x objectives or using an Axioscan microscope (Zeiss) at the 10x and 20x objective. Quantification of paraffin immunohistochemistry and histology was performed using ImageJ (NIH). To determine the number of positively immunostained cells, 4 regions of interest per sample per antibody were manually scored as either positive or negative and reported as percentage positively stained osteoblasts, osteocytes, and chondrocytes per total number of each cell type using ImageJ (NIH). Bone and cartilage area per callus area were calculated using ImageJ (NIH) on H&E and Safranin-O stained sections with 3-4 mice per group per timepoint.

Samples from 7 and 14 dpf were stained with Picro-sirius Red and imaged under polarized light using an Axioscan microscope (Zeiss) at the 20x objective and using second harmonic generated (SHG) microscopy. SHG images were taken on a TCS SP8 Multiphoton Confocal microscope (Leica) at a fundamental wavelength of 880 nm with the 10x and 40x objective on sections oriented in the same direction for all groups. All SHG images were quantified using ImageJ (NIH) and reported as mean pixel intensity within the cortical and callus region relative to WT bone. Mean pixel intensities across four separate regions of interests were averaged as technical replicates for a given sample within either the callus or cortex area with 3-4 mice per group.

Immunofluorescence sections of 4 dpf limbs were imaged on an Axio Observer Z1 (Zeiss) at the 5x, 10x, and 25x objectives. To determine the number of positively EdU-stained periosteal cells, 4 regions of interest were outlined per mouse 1-3 mm from the fracture on the periosteal cortical bone^(28)^. Images were manually scored as either positive or negative, averaged together from each of the 4 regions and reported as percent positively stained per total number of periosteal cells in ImageJ (NIH). Periosteal area and average thickness from these same 4 regions of interested were outlined in ImageJ (NIH) using both immunofluorescence and aqueous H&E sections and averaged together with 6-9 mice per group.

### Periosteal cell isolation and osteogenic differentiation

Mouse periosteal cells were isolated from either WT or Osterix-conditional YAP/TAZ deficient (YAP^cKOi^;TAZ^cKOi^) femurs on 4 dpf and cultured at 37°C and 5% O_2_, as described previously^(29)^. Briefly, mice were anesthetized by carbon dioxide inhalation and euthanized via cervical dislocation. Fractured limbs were carefully dissected of all non-osseous tissue, the epiphyses were then removed, and marrow cavities were flushed. The periosteum was scraped and enzymatically digested for 1 hour at 37°C on an orbital shaker (0.5 mg/ml collagenase P, 2mg/ml hyaluronidase in PBS). Following washing, 2 × 10^4^ cells/cm^2^ were seeded in growth medium (α-MEM, 10% FBS, 1% penicillin-streptomycin, and 1ug/ml doxycycline) and cultured in 5% oxygen for the first 4 days. Half of the media was changed on day 4 and cultures were then incubated in 21% O_2_. Primary cells reached confluence by day 7 and were passaged once into osteogenic differentiation experiments.

Passage 1 periosteal cells were then seeded at 21% O_2_ into 24-well plates (15 × 10^3^ cells/cm^2^) and cultured in growth medium. After reaching confluence, primary periosteal cell cultures were induced towards osteogenic differentiation (50 μg/mL ascorbic acid and 4mM β-glycerophosphate) for 21 days. Osteogenic media was changed every two days prior to RNA isolation and alizarin red staining for mineral deposition at 21 days.

### RNA isolation and qPCR

Limbs from 7 and 14 dpf were carefully dissected and removed of all non-osseous tissue. The intramedullary pin was removed, and marrow flushed before being snap-frozen in liquid nitrogen-cooled isopentane for 1 minute prior to microCT imaging and storage at −80°C until processing. Tissues were then homogenized via mortar and pestle and RNA from the sample was collected using Trizol Reagant (15596026; Life Technologies) followed by centrifugation in chloroform. RNA from fractured limb tissue and cell culture experiments were purified using the RNA Easy Kit (74106; Qiagen) and quantified by spectrophotometry using a NanoDrop 2000 (Thermo-Fisher Scientific). Reverse transcriptase polymerase chain reaction (RT-PCR) was performed on 0.5 µg/µl concentration of RNA using the High-Capacity cDNA Reverse Transcription Kit (4368814; Thermo-Fisher Scientific). Quantitative polymerase chain reaction (qPCR) assessed RNA amount using a StepOnePlus™ Real-Time PCR System (Thermo-Fisher Scientific) relative to the internal control of 18S ribosomal RNA (*18S rRNA*). Data are presented using the ΔΔCt method. Six mice per group were used. Specific mouse primer sequences are listed (**Supplemental Table 1**).

### Statistics and regression

Sample sizes were selected *a priori* by power analyses based on effect sizes and population standard deviations taken from published data on YAP^fl/fl^;TAZ^fl/fl^ mice in other tissues^(30)^, assuming a power of 80% and α=0.05. All statistics and power analyses were performed in GraphPad Prism. Comparisons between two groups were made using the independent t-test while comparisons between 3 or more groups were made using a one-way ANOVA with post-hoc Tukey’s multiple comparisons test, if the data were normally distributed according to D’Agostino-Pearson omnibus normality test and homoscedastic according to Bartlett’s test. When parametric test assumptions were not met, data were log-transformed, and residuals were evaluated. If necessary, either the non-parametric Kruskal-Wallis test with post-hoc Dunn’s multiple comparisons or the non-parametric Mann-Whitney test were used. A p-value < 0.05 (adjusted for multiple comparisons) was considered significant. On the graphs, repeated significance indicator letters (e.g., “a” vs “a”) signify P > 0.05. while groups with distinct indicators (e.g., “a” vs “b”) signify P < 0.05. Summary data are presented as bars, with independent samples indicated in scatter plots and error bars representing standard error of the mean (SEM). Distributions of pMOI were binned and presented as individual samples with lines corresponding to the mean and standard deviation (SD).

## RESULTS

### Constitutive Osterix-conditional YAP and/or TAZ deletion impaired endochondral fracture repair

To determine the roles of YAP and TAZ in endochondral fracture repair, we used Cre-lox to delete YAP and/or TAZ from Osterix-Cre expressing cells prior to embryonic development ^(14)^. We selected a breeding strategy that generated YAP/TAZ allele dosage-dependent Osterix-conditional knockouts with four genotypes (see Table 1). Homozygous YAP/TAZ knockouts were not evaluated due to perinatal lethality ^(16)^. We then evaluated adult bone fracture repair at 14 and 42 days post fracture (dpf).

All genotypes exhibited callus formation by 14 dpf (**Fig. 1A**). However, constitutive YAP and/or TAZ deletion reduced total callus volume and mineralized callus percentage (i.e. BV/TV) at 14 dpf in an allele dosage-dependent manner (**Fig. 1B,C**). Similarly, constitutive YAP and/or TAZ deletion also reduced mineralized tissue volume and volumetric mineral density at 14 dpf (**Supplemental Fig. 1A,B**). We then tested 14 dpf limbs in torsion to failure and observed a similar reduction in maximum torque to failure and torsional rigidity (**Fig. 1D,E**). However, work to max torque and work to failure did not reach statistical significant differences between genotypes (**Supplemental Fig. 1C,D**).

**Figure 1.**
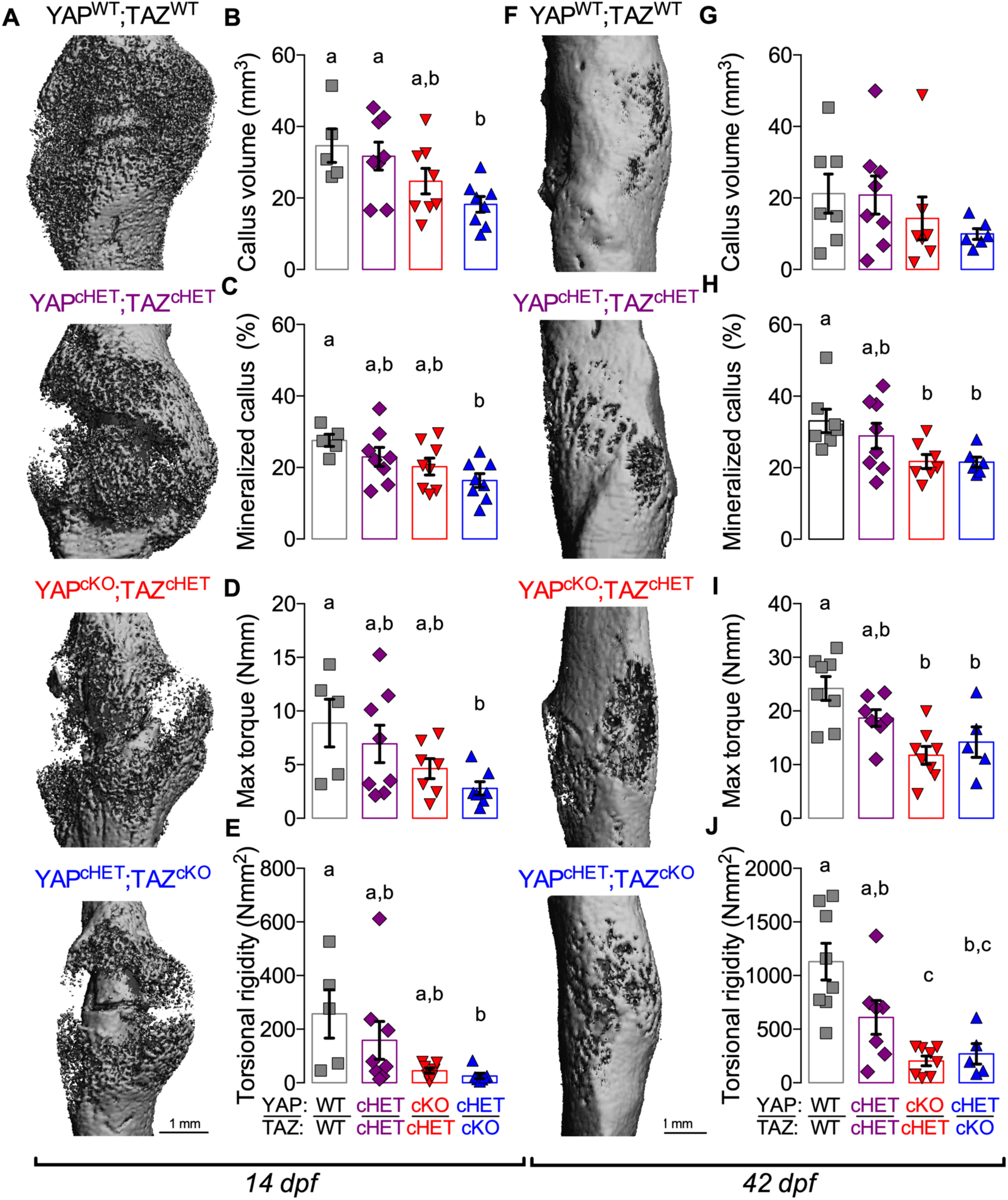
Constitutive, combinatorial YAP/TAZ deletion from Osterix-expressing cells impaired endochondral fracture repair. **A)** MicroCT reconstructions at 14 days post-fracture (dpf). Quantification of 14 dpf callus architecture: **(B)** total callus volume and **(C)** mineralized callus percentage. Quantification of 14 dpf callus mechanical testing in torsion to failure: **(D)** maximum torque and **(E)** torsional rigidity. **F)** MicroCT reconstructions at 42 dpf. Quantification of 42 dpf callus architecture: **(G)** total callus volume and **(H)** mineralized callus percentage. Quantification of 42 dpf callus mechanical testing in torsion to failure: **(I)** maximum torque and **(J)** torsional rigidity. Data are presented as individual samples in scatterplots and bars corresponding to the mean and standard error of the mean (SEM). Sample sizes, N = 5-8. Scale bars indicate 1 mm for microCT reconstructions.

At 42 dpf, all genotypes similarly underwent hard callus remodeling (**Fig. 1F**). However, constitutive YAP and/or TAZ deletion again reduced mineralized callus percentage, maximum torque to failure, and torsional rigidity, but at this timepoint differences in total callus volume did not reach statistical significance (**Fig 1G-J**). At 42 dpf, constitutive YAP and/or TAZ deletion reduced volumetric mineral density, but did not significantly reduce mineralized tissue volume, work to maximum torque, or work to failure (**Supplemental Fig. 1E-H**).

### Constitutive Osterix-conditional YAP and/or TAZ deletion reduced callus formation

To determine the distribution of the callus, including cartilage and bone, at 14 dpf, we quantified total callus polar moment of inertia (**Fig. 2A**). Independent of bone formation, constitutive YAP/TAZ deletion reduced cartilage callus formation, particularly in the YAP^cHET^;TAZ^cKO^ mice, which are homozygous for TAZ deletion and heterozygous for YAP (**Fig. 2B**). At 42 dpf, similar results were observed where the polar moment of inertia distribution of mineralized tissue within the callus were reduced in the YAP^cHET^;TAZ^cKO^ mice (**Supplemental Fig. 2A,B**). At 42 dpf, we performed an analysis of covariance (ANCOVA) using linear regression to decouple the contributions of callus mineralization and geometric distribution from the mechanical behavior^(16,31)^, since constitutive YAP/TAZ deletion reduced both callus mineralization and geometry. We found that individual regression lines for each genotype best predicted maximum torque to failure and torsional rigidity, suggesting that differences in connectivity or composition also contribute to mechanical behavior (**Supplemental Fig. 2C,D**).

**Figure 2.**
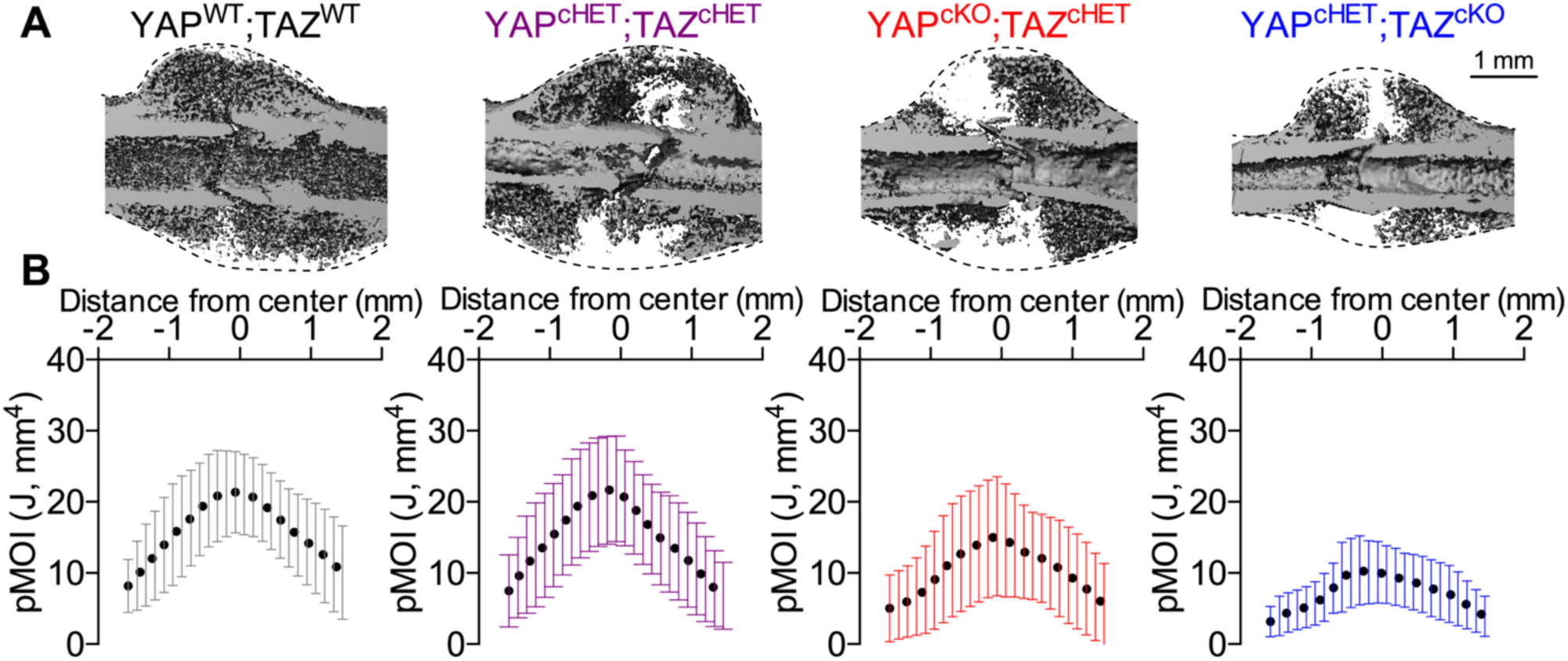
Constitutive, combinatorial YAP/TAZ deletion from Osterix-expressing cells reduced overall callus size. **A)** MicroCT reconstructions at 14 dpf showing longitudinal cut-planes within the callus. Dotted lines indicate the callus boundary, the region in which moment of inertia was quantified. **B)** Polar moment of inertia distributions of the entire callus for each of the YAP/TAZ allele dose-dependent knockout genotypes. Data were binned into 25 equal distance bins from the center of the callus and presented as dots representing the mean and error bars corresponding to the standard deviation (SD). Sample sizes, N = 5-8. Scale bar indicates 1 mm for microCT reconstructions.

### Constitutive Osterix-conditional YAP and/or TAZ deletion impaired periosteal development

To determine whether developmental defects in the periosteum contributed to the reduced callus formation, we evaluated the periosteal thickness and cellularity of intact femurs from constitutive knockout mice. Constitutive YAP and/or TAZ deletion significantly reduced periosteal thickness (**Fig. 3A-C**). Similar to younger mice^(16)^, constitutive YAP and/or TAZ deletion reduced periosteal cell number per bone surface in an allele dose-dependent manner (**Fig 3. D**), suggesting defective periosteal development.

**Figure. 3.**
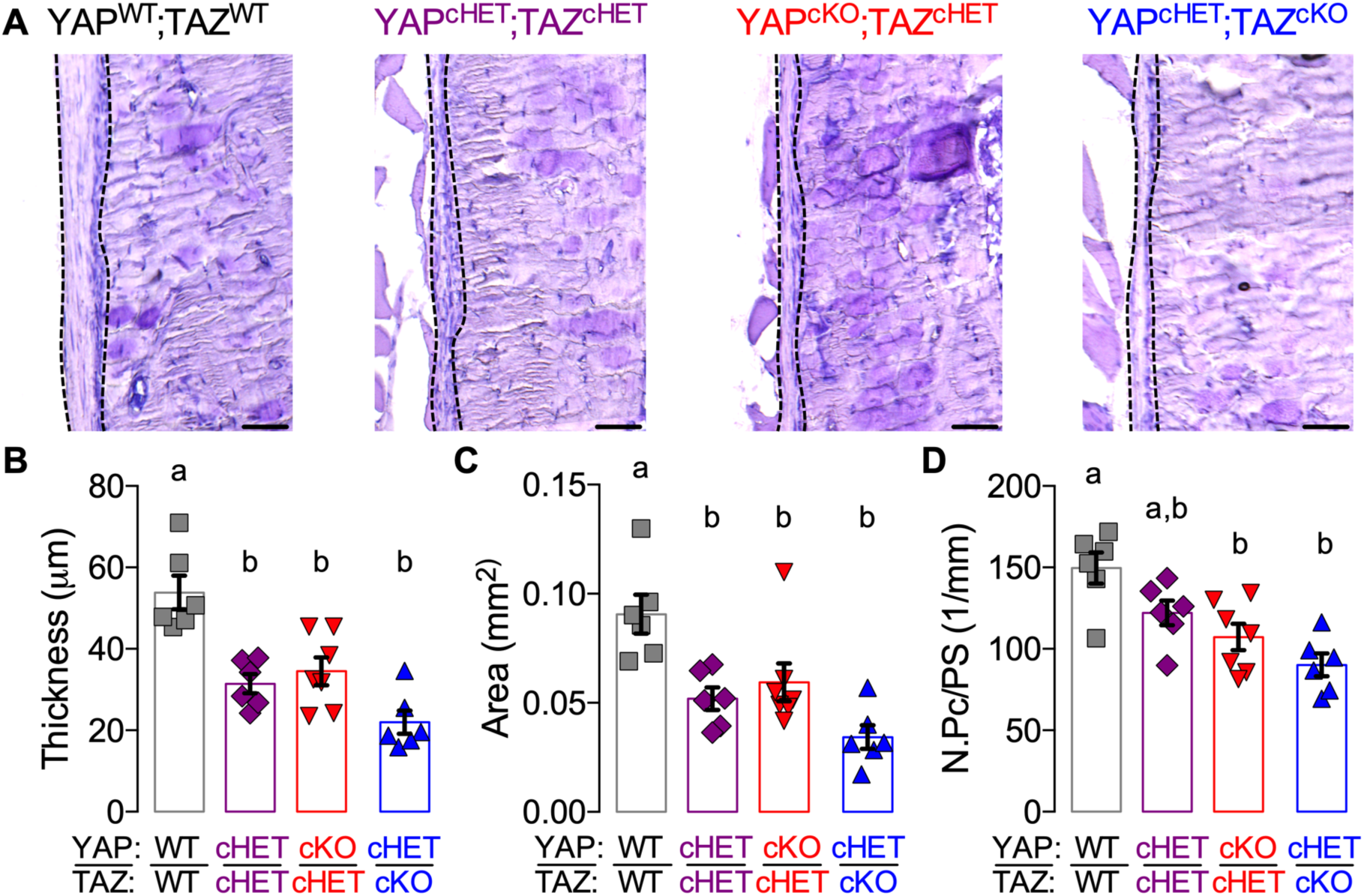
Constitutive, combinatorial YAP/TAZ deletion from Osterix-expressing cells impaired periosteal development in intact bone. **A)** Representative micrographs of 18-21 weeks-old distal femur cortical bone stained by aqueous H&E. Dotted lines indicate the periosteum. Quantification of **(B)** periosteal thickness, **(C)** periosteal area and **(D)** periosteal cell number per bone surface (N.Pc /PS). Data are presented as individual samples in scatterplots with bars corresponding to the mean and standard error of the mean (SEM). Sample sizes N = 6-7. Scale bars indicate 50 μm for all images.

### Inducible Osterix-conditional YAP/TAZ deletion impaired callus mineralization, but not formation

The impairment of endochondral bone fracture healing observed in the constitutive YAP/TAZ deletion knockout model could be a consequence of defective periosteal stem cell supply, expansion, and/or differentiation. To address this question, we generated adult-inducible, dual homozygous YAP/TAZ knockout mice (YAP^cKOi^;TAZ^cKOi^) in which the periosteal progenitor population was allowed to develop normally prior to fracture. Here, we induced homozygous Osterix-conditional YAP/TAZ deletion two weeks prior to fracture. During those two weeks prior to fracture, Osterix-conditional YAP/TAZ deletion did not significantly affect periosteal cell thickness, area, or cell number (**Fig. 4A-D**).

**Figure 4.**
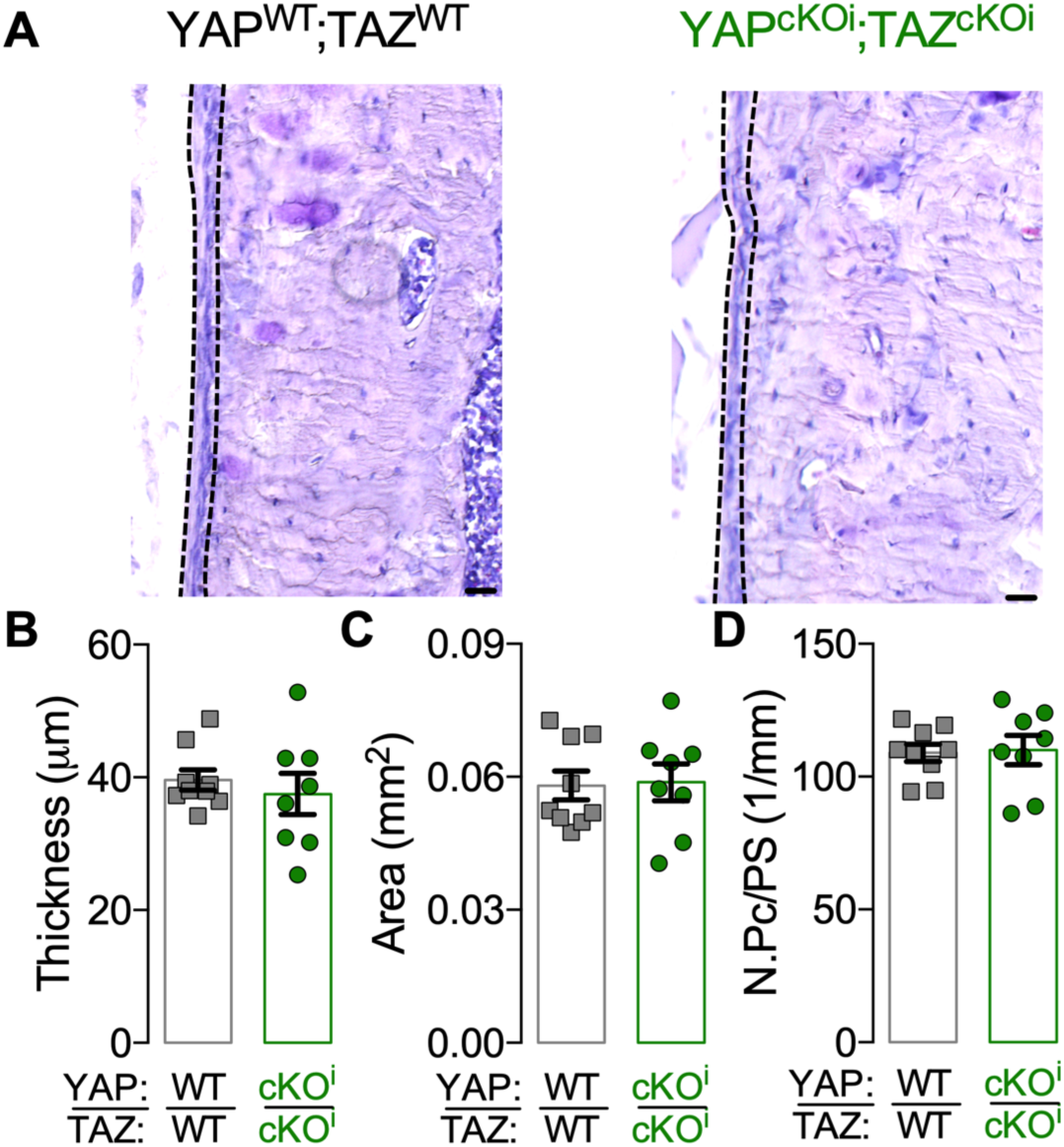
Periosteal thickness and cellularity developed normally in adult onset-induced, Osterix-conditional, homozygous YAP/TAZ knockout mice. **A)** Representative micrographs of 16-18 weeks-old cortical bone stained by aqueous H&E. Dotted lines indicate the periosteum. Quantification of **(B)** periosteal thickness, **(C)** periosteal area and **(D)** periosteal cell number per bone surface (N.Pc/PS). Data are presented as individual samples in scatterplots and bars corresponding to the mean and standard error of the mean (SEM). Sample sizes N = 8-9. Scale bars indicate 30 μm for all images.

To analyze the recombination efficiency of inducible Osterix-Cre-mediated YAP/TAZ deletion following fracture, we evaluated YAP/TAZ expression in chondrocytes, osteoblasts, and osteocytes within the callus at 14 dpf by immunohistochemistry and qPCR (**Supplemental Fig. 3A-C**). YAP protein expression was significantly reduced in chondrocytes (23% reduction), osteoblasts (23% reduction), and osteocytes (26% reduction) (**Supplemental Fig. 3D-F**). YAP mRNA expression was reduced by 59% in YAP^cKOi^;TAZ^cKOi^ callus lysate preparations (**Supplemental Fig. 3G**). TAZ protein expression was moderately reduced in chondrocytes (11% reduction; p = 0.1) and significantly reduced in osteoblasts (18% reduction), and osteocytes (12% reduction) (**Supplemental Fig. 3H-J**). TAZ mRNA expression was reduced by 49% in YAP^cKOi^;TAZ^cKOi^ callus lysate preparations (**Supplemental Fig. 3K**).

All genotypes underwent initial callus formation by 14 dpf (**Fig. 5A**). Inducible Osterix-Cre-mediated YAP/TAZ deletion reduced mineralized callus percentage and volumetric bone mineral density, but differences in total callus volume between groups were not observed (**Fig. 5B-D**). Further, inducible Osterix-Cre-mediated YAP/TAZ deletion significantly increased variability in total callus volume size and the polar moment of inertia distribution within the callus in comparison to YAP^WT^;TAZ^WT^ and Osx:Cre mice (**Fig. 5B; Supplemental Fig. 4A,B**).

**Figure 5.**
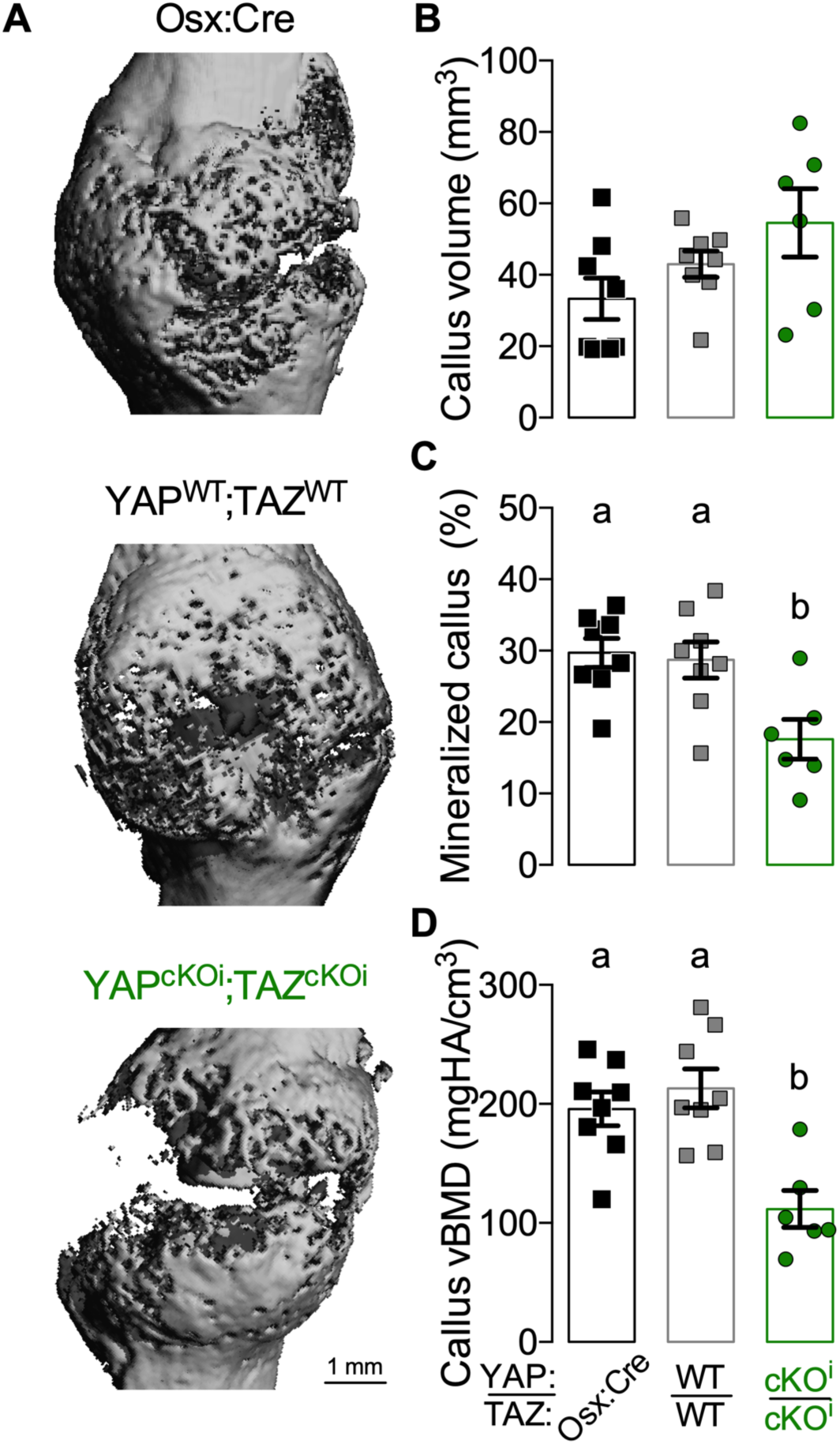
Adult onset-inducible, homozygous YAP/TAZ deletion from Osterix-expressing cells impaired callus mineralization, but not formation. **A)** MicroCT reconstructions at 14 dpf. Quantification of 14 dpf callus architecture: **(B)** total callus volume, **(C)** mineralized callus percentage, and **(D)** volumetric mineral density. Data are presented as individual samples in scatterplots and bars corresponding to the mean and standard error of the mean (SEM). Sample sizes, N = 6-8. Scale bars indicate 1 mm for microCT reconstructions.

### Inducible Osterix-conditional YAP/TAZ deletion did not alter cartilaginous callus formation, but reduced bone formation

As the formation of a cartilaginous callus template is a critical step during endochondral fracture healing^(3–5)^, we histologically evaluated cartilage formation at 14 dpf (**Supplemental Fig. 5A,B**). Consistent with microCT, inducible Osterix-Cre-mediated YAP/TAZ deletion did not affect total callus area at 14 dpf, but increased variability was observed YAP^cKOi^;TAZ^cKOi^ mice (**Supplemental Fig. 5C)**. Differences in total cartilage area and percent cartilage area were not detected between groups (**Supplemental Fig. 5D,E)**. However, inducible Osterix-Cre-mediated YAP/TAZ deletion resulted in a non-significant trend towards increase percent cartilage area at 14 dpf (p = 0.12; **Supplemental Fig. 5E**). At 14 dpf, inducible Osterix-Cre-mediated YAP/TAZ deletion did not significantly alter mRNA expression of markers for chondrogenesis, including SRY-Box Transcription Factor 9 (*Sox9*) and Aggrecan (*Acan*), but resulted in a non-significant trend towards reduced collagen, type II, alpha 1 (*Col2a1*) mRNA expression (p = 0.06; **Supplemental Fig. 5F**).

Following formation of the cartilaginous callus, extensive bone formation occurs within the callus through both intramembranous and endochondral ossification^(4)^. Thus, we next histologically evaluated endochondral ossification and bone formation at 14 dpf (**Fig. 6A,B**). At 14 dpf, differences in Osterix (OSX)-positive hypertrophic chondrocytes were not detected between groups (**Fig. 6C)**. At 14 dpf, the number of OSX-positive osteoblasts per bone surface, bone area, and percent bone area were reduced following inducible Osterix-Cre-mediated YAP/TAZ deletion (**Fig. 6D-F)**. Further, inducible Osterix-Cre-mediated YAP/TAZ deletion reduced mRNA expression of markers for hypertrophic chondrocytes, including collagen, type X, alpha 1 (*Col10*) and vascular endothelial growth factor (*Vegfa)* (**Fig.6J**). Similarly, inducible Osterix-Cre-mediated YAP/TAZ deletion reduced mRNA expression of markers of collagen, including collagen type I, alpha 1 (*Col1a1*) and collagen type I, alpha II (*Col1a2*), but not serpin family H member 1 (*SerpinH1*) (**Fig.6K**). Lastly, inducible Osterix-Cre-mediated YAP/TAZ deletion reduced mRNA expression markers of osteogenesis, including osteoblast-specific transcription factor Osterix (*Osx*) and alkaline phosphatase (*Alp*), while reductions in runt-related transcription factor 2 (*Runx2*) or bone sialoprotein (*Bsp*) did not reach statistical significance (**Fig.6L)**.

**Figure. 6.**
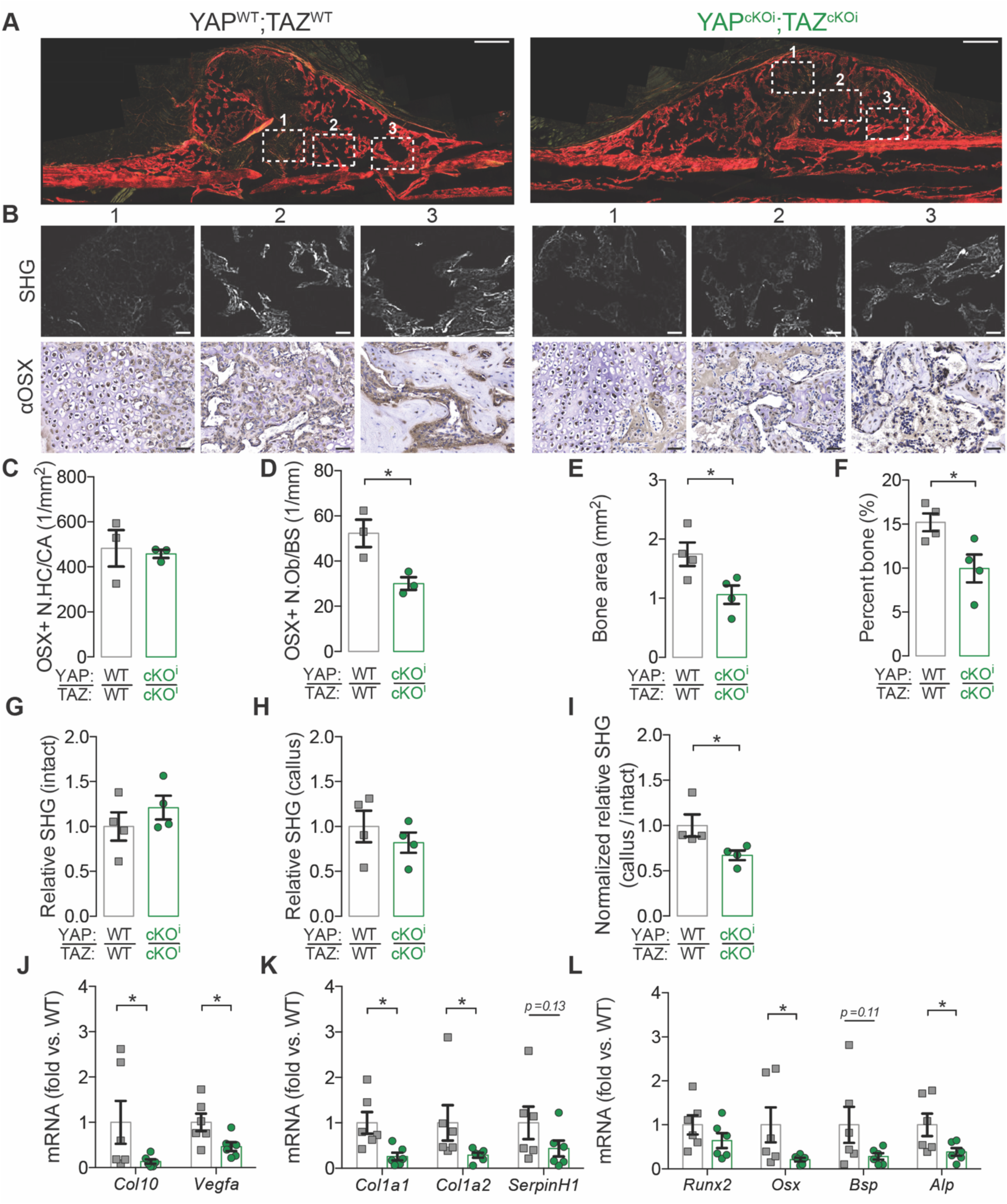
Adult onset-inducible, homozygous YAP/TAZ deletion from Osterix-expressing cells reduced callus bone formation. **A)** Micrographs of Picrosirius Red-stained calluses at 14 dpf. White boxes outline three regions of interest **(B)** in which second harmonic generated imaging (SHG) and anti-Osterix (αOSX) immunostaining are highlighted. Quantification of callus histomorphometry at 14 dpf of **(C)** the number Osterix-positive hypertrophic chondrocytes per cartilage area (OSX+ N.HC/CA), **(D)** Osterix-positive osteoblasts per bone surface (OSX+ N.Ob/BS), **(E)** bone area, and **(F)** percent bone area. Quantification of relative SHG intensity per bone area at 14 dpf of **(G)** the intact cortical bone, **(H)** the newly formed callus bone, and **(I)** normalized callus-to-intact SHG intensity. Messenger RNA was extracted from callus lysate preparations at 14 dpf and target gene expression was normalized to *18S rRNA* and quantified as fold-change relative to wild type (**J-L**). **J)** *Col10* and *Vegfa* mRNA expression. **K)** *Col1a1, Col1a2*, and *SerpinH1* mRNA expression. **L)** *Runx2, Osx, Bsp*, and *Alp* mRNA expression. Data are presented as individual samples in scatterplots and bars corresponding to the mean and standard error of the mean (SEM). N = 6 per group for qPCR and N = 3-4 per group for histomorphometry. Scale bars indicate 50 μm for all high-power images and 500 μm for callus images.

### Inducible Osterix-conditional YAP/TAZ deletion reduced periosteal osteoblast precursor expansion in vivo and osteogenic differentiation in vitro

Given the defect in osteogenesis resulting from inducible Osterix-Cre-mediated YAP/TAZ deletion, we evaluated activated periosteal progenitors at 4 dpf in four regions distal and proximal to the fracture line, in which periosteal osteoblast precursors are primarily fated to form bone through direction intramembranous ossification^(32)^ (**Fig. 7A)**. At 4 dpf, inducible YAP/TAZ deletion reduced periosteal osteoprogenitor cell expansion in terms of total area and average thickness (**Fig. 7B-D**). However, inducible YAP/TAZ deletion did not reduce the number of periosteal osteoprogenitor cells per expanded periosteal area, but significantly reduced the percentage of proliferating, EdU-positive periosteal osteoprogenitor cells (**Fig. 7E,F**).

**Figure 7.**
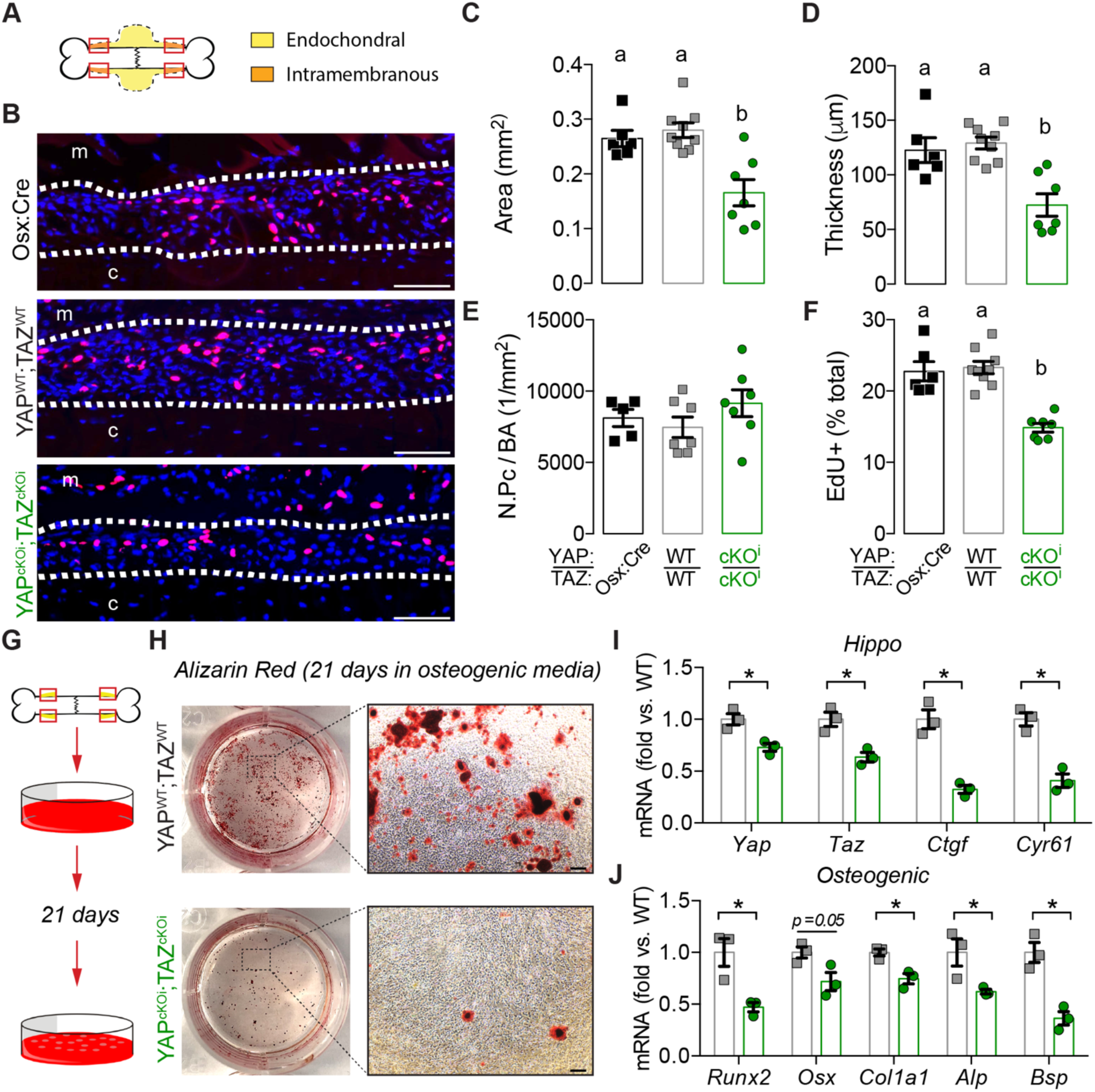
Adult onset-inducible, homozygous YAP/TAZ deletion from Osterix-expressing cells impaired periosteal osteoblast precursor cell expansion and osteogenic differentiation. **A)** Representation of the regions of interest in the callus “shoulder,” where bone formation initiates by intramembranous ossification, and where we evaluated periosteal osteoblast precursor expansion and proliferation. **B)** Representative micrographs of EdU+ periosteal cells (red) at 4 dpf, with all nuclei counterstained by DAPI (blue). White dotted lines indicate periosteal cell expansion zone; “m” indicates muscle, “c” indicates cortical bone. Quantification of the expanded **(C)** periosteal area, **(D)** average thickness, **(E)** number of periosteal cells per bone area (N.Pc/BA), and **(F)** percentage of EdU-positive periosteal cells. **G)** Activated periosteal cells isolated from 4 dpf limbs were cultured in osteogenic media for 21 days. **H)** Representative Alizarin Red staining of mineral deposition following osteogenic induction. **I)** *Yap, Taz, Ctgf*, and *Cyr61* and **J)** *Runx2, Osx, Alp*, and *Bsp* mRNA expression, relative to *18S rRNA*, from periosteal progenitor cell cultures following 21 days of osteogenic induction. Data are presented as individual samples in scatterplots and bars corresponding to the mean and standard error of the mean (SEM). N = 6-9 per group. Scale bars indicate 100 μm for all high-power EdU images and 1 mm for high-power Alizarin Red images.

Though unexplored in periosteal osteoprogenitor cells, YAP and TAZ are known to mediate osteogenic differentiation in the MSCs^(33–37)^, which originate from a common mesenchymal embryonic lineage^(38)^. To elucidate if inducible YAP/TAZ deletion regulated periosteal osteoprogenitor differentiation, we isolated activated periosteal progenitor cells at 4 dpf (**Fig. 7G**)^(28,29)^. Following culture for 21 days in osteogenic media, inducible, Osterix-conditional YAP/TAZ deletion reduced mineral deposition stained with Alizarin Red (**Fig. 7G-H**). As expected, Osterix-conditional inducible YAP/TAZ deletion *in vitro* reduced mRNA expression of *Yap* and *Taz* as well as their canonical downstream target, *Ctgf* and *Cyr61* in periosteal progenitor cells from YAP^cKOi^;TAZ^cKOi^ mice (**Fig. 7I**). Lastly, Osterix-conditional inducible YAP/TAZ deletion *in vitro* reduced mRNA expression of osteogenic differentiation genes including, *Runx2, Col1a1, Alp, and Bsp* while reductions in *Osx* (p = 0.05) did not reach statistical significance (**Fig. 7H**).

Given the defective periosteal osteoblast precursor expansion and osteogenic differentiation, we evaluated bone formation in the callus at 7 dpf. Although all genotypes underwent periosteal expansion and mineralization (**Fig. 8A**), inducible Osterix-Cre-mediated YAP/TAZ deletion reduced mineralized callus percentage and volumetric mineral density at 7 dpf (**Fig. 8B,C**). Within regions of the callus undergoing intramembranous bone formation (**Fig. 8D**), inducible Osterix-Cre-mediated YAP/TAZ deletion reduced OSX-positive osteoblasts per bone surface concomitant with reduced bone percent area while resulting in a non-significant trend toward reduced bone area (**Fig. 8E-H**), consistent with our observations at 4 dpf *in vivo* and *in vitro*. At 7 dpf, significant changes in cartilaginous callus formation were not detected (**Supplemental Fig. 6A-F)**. Similarly, inducible Osterix-Cre-mediated YAP/TAZ deletion did not significant alter endochondral ossification or matrix collagen content and organization within the callus (**Supplemental Fig. 7A-I)**.

**Figure 8.**
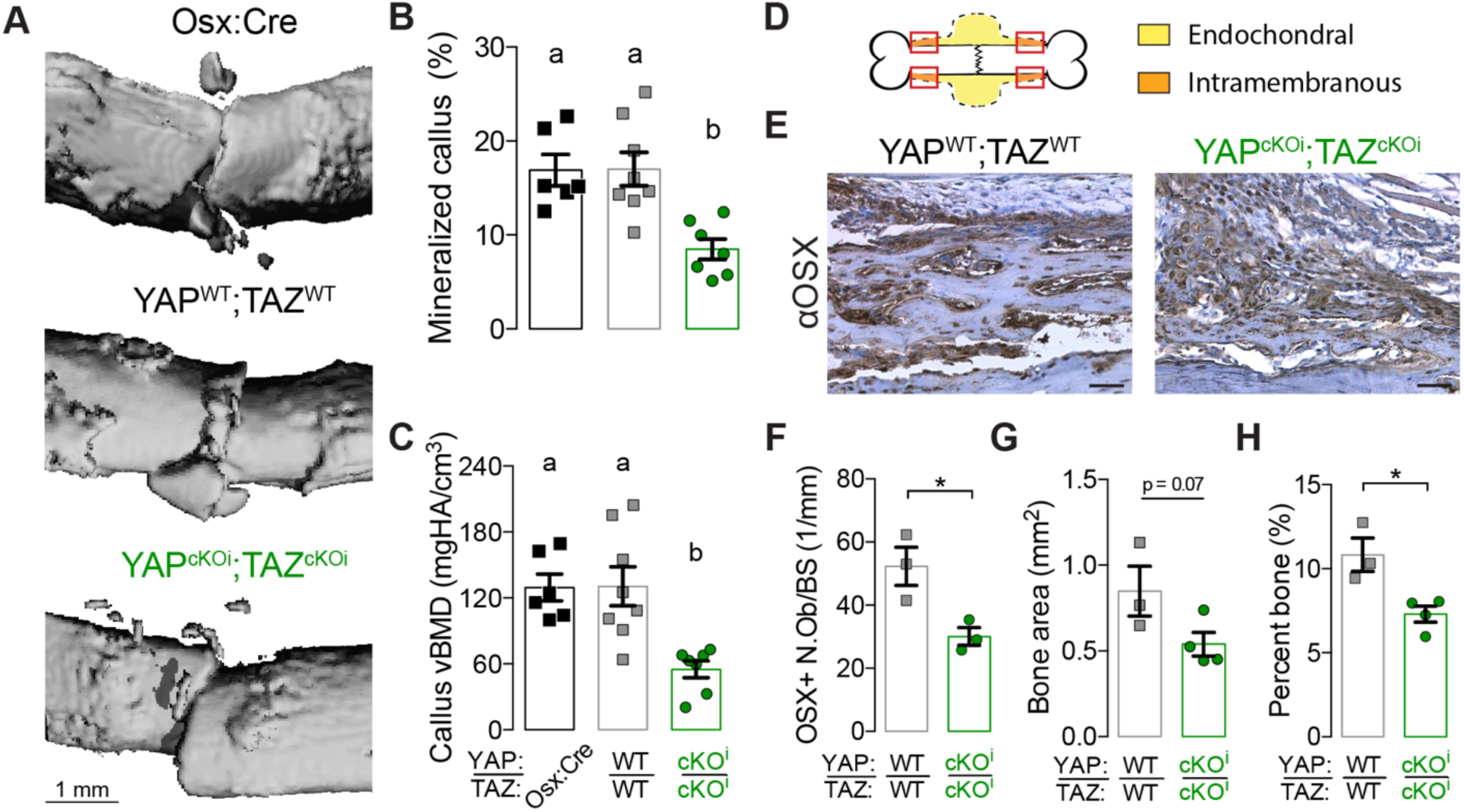
Adult onset-inducible, homozygous YAP/TAZ deletion from Osterix-expressing cells impaired periosteal osteoblast precursor bone formation. **A)** MicroCT reconstructions at 7 days post-fracture (dpf). Quantification of 7 dpf callus architecture: **(B)** mineralized callus percentage and **(C)** volumetric bone mineral density. **D)** Representation of the regions of interest in the callus “shoulder,” where bone formation initiates by intramembranous ossification, and where we evaluated (**E**) anti-Osterix (αOSX) immunostaining at 7 dpf. Quantification of callus histomorphometry at 7 dpf of **(F)** Osterix-positive osteoblasts per bone surface (OSX+ N.Ob/BS). **(G)** bone area, and **(H)** percent bone area. Data are presented as individual samples in scatterplots and bars corresponding to the mean and standard error of the mean (SEM). Sample sizes, N = 6-8. Scale bars indicate 1 mm for microCT reconstructions and 50 μm for micrographs.

## DISCUSSION

This study identifies new roles for the transcriptional regulators, YAP and TAZ, in bone fracture healing, adding to our understanding of periosteal osteoblast precursor cell regulation. Here, we show that YAP and TAZ promote expansion and osteoblastic differentiation of periosteal osteoblast precursors to promote bone fracture healing. Constitutive YAP and/or TAZ deletion from Osterix-expressing cells impaired bone fracture healing by reducing both callus formation and subsequent mineralization, due in part developmental defects in the periosteal progenitor supply. When we allowed for the development of a normal periosteal progenitor population prior to fracture, adult onset-induced YAP/TAZ deletion did not impair cartilaginous callus formation, but delayed mineralization due to impaired osteoblast precursor cells and their osteogenic differentiation. Together, these data demonstrate that the transcriptional co-activators, YAP and TAZ, promote the expansion and differentiation of periosteal osteoblast precursors to accelerate bone fracture healing.

Fracture healing recapitulates many aspects of embryonic skeletal development, but features a unique post-natal environment, resulting in contextual differences^(3,39)^. We previously found that Osterix-conditional YAP/TAZ deletion in the embryo caused a severe skeletal fragility phenotype^(16)^, while Xiong et al. induced Osterix-conditional YAP/TAZ deletion at post-natal day 21 (P21) and performed skeletal phenotyping at P84, observing increased osteoblast numbers but no measurable effect on whole bone microarchitecture^(17)^. Our present data resolve the differences between these two studies, establishing a critical role for YAP and TAZ in the development of the postnatal osteoprogenitor niche and demonstrating critical roles for YAP and :TAZ in osteoblast precursor proliferation and differentiation in a context of rapid bone formation, similar to that which occurs during bone development, in contrast to postnatal growth and homeostasis. The present data are further consistent with other reports. For example, deletion of the YAP/TAZ-regulated transcription factors, Snail and Slug, from Osterix-expressing cells reduced both the proliferative potential of bone surface-associated osteoprogenitors and osteogenic differentiation capacity of adult skeletal stem cells^(40)^. Similarly, conditional deletion of YAP from Osteocalcin-expressing cells reduced osteoblast progenitor cell proliferation and differentiation, further supporting a role for YAP and TAZ in promoting osteoblast progenitor cell function ^(18)^.

Endochondral bone fracture repair includes both the formation of a cartilage template as well as subsequent osteoblast-mediated mineralization^(3,41)^. Here, we found that while constitutive deletion impaired callus formation, adult onset-inducible YAP/TAZ deletion did not significantly affect cartilage formation during fracture healing. A previous study found that YAP overexpression in developing chondrocytes, using the Col2a1 promoter, as well as deletion of MST1/2 using Dermo-Cre impaired endochondral fracture healing^(42)^, which appears to contradict the results described here. However, both models exhibit a developmental skeletal phenotype prior to fracture^(42)^, and in particular, observed that YAP/TAZ negatively regulate chondrogenesis. Thus, inducible targeting models are needed to decouple the developmental history from the process of fracture repair ^(43,44)^. Here, we selected the tetOFF Osterix-Cre inducible system instead of the tamoxifen-inducible Osterix-Cre^ERT2 (45)^, as tamoxifen is rapidly cleared^(46)^, resulting in transient Cre-activity^(47)^ in newly-generated Osterix-positive cells during fracture repair. Furthermore, targeting conditional gene inactivation in chondrocytes versus osteoblast-lineage cells can result in drastically different phenotypes. For example, conditional deletion of Runx2 in chondrocytes using Col2a1-Cre phenocopied global Runx2 gene inactivation with perinatal lethality and a lack of mineralization while conditional deletion of Runx2 in osteoblasts using 2.3kb-Col1a1-Cre resulted in a moderate low bone mass phenotype^(48,49)^. Nonetheless, hypertrophic chondrocytes are known to express Osterix during endochondral ossification^(50–52)^ and we observed moderate Osterix-Cre-mediated YAP/TAZ recombination in hypertrophic chondrocytes, suggesting that the relative contributions of YAP and TAZ during endochondral ossification are potentially stage-dependent^(16–18,42)^. Future studies will identify the temporal and cell-specific contributions of YAP and TAZ to bone development and repair.

Recent and ongoing studies have revealed remarkable diversity in both the cellular identity and regulatory signals that contribute to periosteal function. Gli1^(53)^, Prx1^(38)^, αSMA^(54)^, cathepsin K^(55)^, and Osterix^(56)^ mark both overlapping and distinct periosteal progenitor cell populations, while markers previously thought to define bone marrow stromal cells, including CD73, CD90, CD105, PDGFRα, Gremlin 1, Cxcl12, and Nestin^(38,57–60)^, also show high expression in the periosteal progenitors. Ongoing efforts continue to uncover new skeletal progenitor cell populations that contribute to fracture repair^(59,61–65)^, and the intersection of this cellular diversity with YAP/TAZ signaling remains unclear. Here, we observed significant reductions in mRNA expression of endochondral-, collagen-, and osteogenesis-related gene expression signatures at 14 dpf, which can be explained either by indirect shifts in the cell populations that express these targets or by direct YAP/TAZ-mediated transcriptional regulation of those genes. Further research will be required to not only systematically identify the transcriptional co-effectors of YAP and TAZ in each cell type of interest, but also delineate the periosteal progenitor subpopulations affected by YAP/TAZ signaling.

YAP and TAZ may regulate osteoblast precursor cell proliferation and osteogenic differentiation through a variety of mechanisms. Conditional deletion of YAP in osteoblasts using Osteocalcin-Cre reduced osteoblast progenitor proliferation as well as osteogenic differentiation and proposed YAP stabilized β-catenin to promote β-catenin-mediated osteogenesis^(18)^. However, evidence exists for YAP and TAZ playing both a positive and negative role in WNT/β-catenin signaling^(35,66)^, suggesting that further investigating into YAP/TAZ-dependent regulation of this pathway in periosteal progenitors is needed. A similar study demonstrated that Snail and Slug form stable protein-protein complexes with both YAP and TAZ in tandem to promote osteoprogenitor proliferation and differentiation^(40)^. In osteoprogenitors, the Snail/Slug-YAP/TAZ axis promotes proliferation by interacting with TEAD to enhance TEAD-dependent transcriptional activity and downstream expression of YAP/TAZ-TEAD target genes, such as *Ctgf* and *Ankrd1*^(40)^. In contrast, the Snail/Slug-YAP/TAZ axis promotes osteogenic differentiation via Snail/Slug-TAZ interactions with Runx2 to promote Runx2-dependent transcriptional activity and downstream expression of osteogenic target genes, such as *Osterix* and *Alp*^(40)^. Accordingly, evidence for TAZ interacting with Runx2 to promote downstream osteogenic gene expression *in vitro* is strong ^(37,67)^, but YAP has been observed to both inhibit and promote downstream osteogenic gene expression *in vitro*^(68–72)^. Thus, future studies to identify the molecular mechanisms by which YAP and TAZ control periosteal progenitor expansion and differentiation are needed.

This study has several limitations. First, small sample sizes (N =3-9) produced under-powered analyses for some outcome measures. Thus, not all the studies performed had sufficient power to compare sex as a variable, though we did not observe sexually dimorphic behavior for any outcome measure, and our prior assessment of YAP/TAZ regulation of bone development did not show an effect of sex^(16)^. Second, we used an open fracture model, which may affect the kinetics and immunology of the fracture repair process^(73,74)^. We initially began these experiments using a closed fracture model, following the Einhorn method^(75)^, but this produced a high percentage of comminuted fractures in the YAP/TAZ cKO groups. We therefore moved to an open fracture model in which the bending moment could be applied with lower kinetic energy. This observation suggests that YAP/TAZ deletion during development impaired the bone matrix fracture toughness, consistent with our prior report on the bone fragility phenotype^(16)^. Third, adult onset-inducible Osterix-conditional knockout increased variability in response to fracture, adding an additional layer of complexity to the already challenging study of endochondral bone fracture healing biology^(5)^. The drug used to prevent Cre-mediated recombination, doxycycline, is a tetracycline derivative. Tetracycline exhibits high affinity for exposed mineral and is therefore commonly used as a label for dynamic bone histomorphometry^(76)^. Potential embedding of doxycycline into the bone matrix during skeletal development and subsequent release following fracture could impair robust Cre-recombination and reduce the observed effect size for adult onset-inducible YAP/TAZ knockout mice. We recommend additional study to quantify the kinetics of tetOff inducible systems and efficiency of Cre-mediated inducible recombination in bone. Lastly, the Osterix-Cre transgene is known to cause defects in craniofacial development^(77)^ and in fracture callus formation^(52)^, depending on genetic background. However, on this background, we did not observe differences between Osx-Cre and YAP^WT^;TAZ^WT^ wild type mice, demonstrating phenotypic specificity for YAP/TAZ deletion.

In conclusion, this study identifies the transcriptional co-activators, YAP and TAZ, as regulators of bone fracture healing that promote periosteal osteoblast precursor proliferation and osteogenic differentiation to accelerate bone healing. Further elucidation of the mechanisms by which YAP and TAZ control the periosteal progenitor cell response to fracture may help guide the development of future targeted therapies to enhance bone fracture healing.

## ACKNOWLEDGEMENTS

YAP^fl/fl^;TAZ^fl/fl^ mice were provided by Dr. Eric Olson (University of Texas Southwestern Medical Center). Theresa Sikorski (University of Notre Dame) provided initial mouse husbandry and maintenance. The authors declare no conflicts of interest. C.D.K., L.Q. and J.D.B. designed research; C.D.K., M.P.N., Y.M., and J.D.B. analyzed data; C.D.K., M.P.N., Y.M., H.B.P., J.H.D., and K.M.J. performed research; L.Q. contributed new reagents/analytic tools; C.D.K. and J.D.B. wrote the paper; All authors reviewed and approved the paper. C.D.K and J.D.B. take responsibility for integrity of data analysis. This work was supported by the National Institute of Arthritis, Musculoskeletal, and Skin Diseases (NIAMS) of the National Institutes of Health through grants T32 AR007132 (to C.D.K.), P30 AR069619 (to J.D.B.), and R01 AR074948 (to J.D.B.).

## Notes

**GRANT SUPPORTERS**: This work was supported by the National Institute of Arthritis, Musculoskeletal, and Skin Diseases (NIAMS) of the National Institutes of Health through grants T32 AR007132 (to C.D.K.), P30 AR069619 (to J.D.B.), and R01 AR074948 (to J.D.B.).

**DISCLOSURES**: The authors have nothing to disclose.

