## Supplemental Data for "YAP and TAZ promote periosteal osteoblast precursor expansion and differentiation for fracture repair"

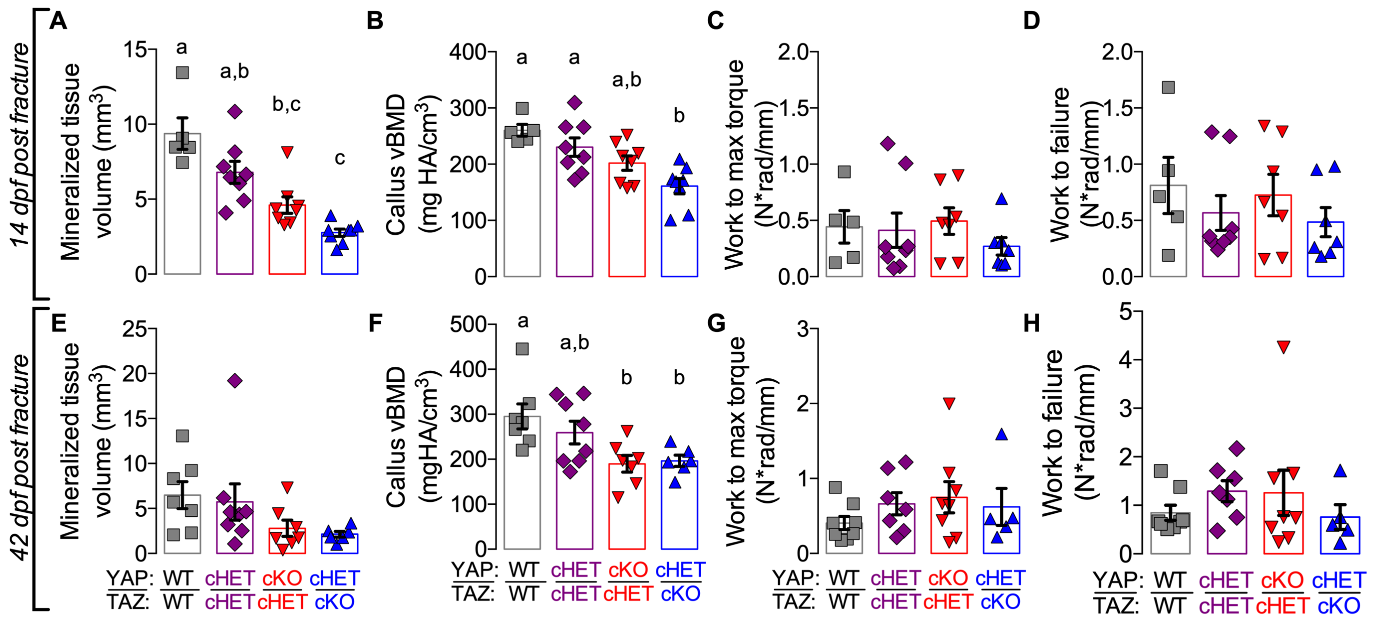


**Supplemental Fig. 1. Constitutive, combinatorial YAP/TAZ deletion from Osterix-expressing cells impaired endochondral fracture repair, but not callus toughness.** 14 and 42 dpf calluses were analyzed using microCT and torsion testing to failure. Quantification of 14 dpf callus architecture: **(A)** total mineralized tissue volume and **(B)** volumetric mineral density. Quantification of 14 dpf callus mechanical testing in torsion to failure: **(C)** work to maximum torque and **(D)** work to failure. Quantification of 42 dpf callus architecture: **(E)** total mineralized tissue volume and **(F)** volumetric mineral density. Quantification of 42 dpf callus mechanical testing in torsion to failure: **(G)** work to maximum torque and **(H)** work to failure. Data are presented as individual samples in scatterplots and bars corresponding to the mean and standard error of the mean (SEM). Sample sizes, N = 5-8.

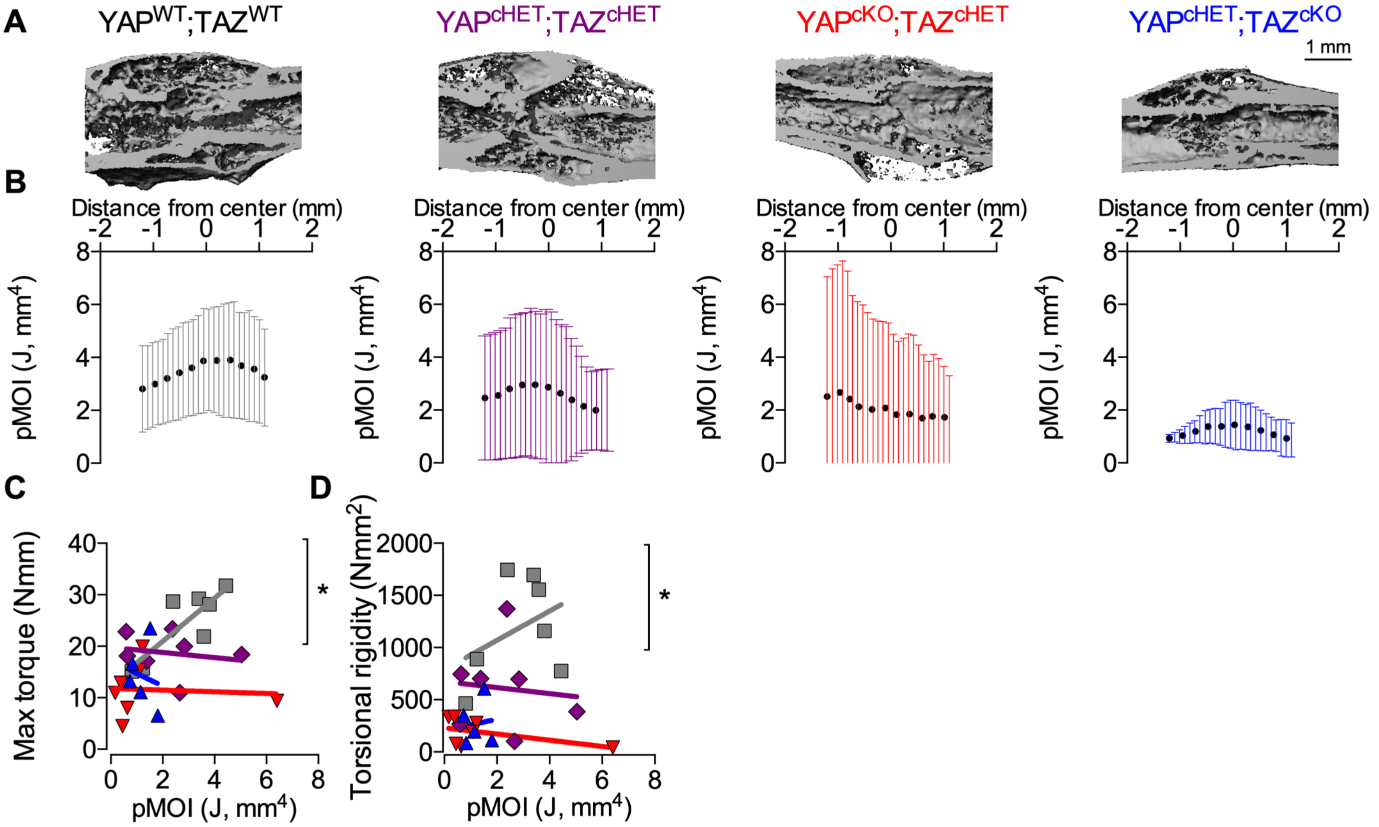


**Supplemental Fig. 2. Combinatorial, constitutive YAP/TAZ deletion from Osterix-expressing cells reduced hard callus size and mechanical properties. A)** 42 dpf post fracture microCT reconstruction of longitudinal cut planes within the callus. Mineralized tissue within the callus were included in polar moment of inertia analysis **B)** Polar moment of inertia distributions of the mineralized limb for each of the YAP/TAZ allele dose dependent knockout genotypes. Data were binned into 25 equal distance bins from the center of the callus and presented as dots representing the mean and bars corresponding to the standard deviation (SD). ANCOVA analysis accounting for fractured limb geometry revealed significant differences between genotypes in **(C)** failure properties and **(D)** elastic properties. Sample sizes, N = 5-8. Scale bars indicate 1 mm for microCT reconstructions.

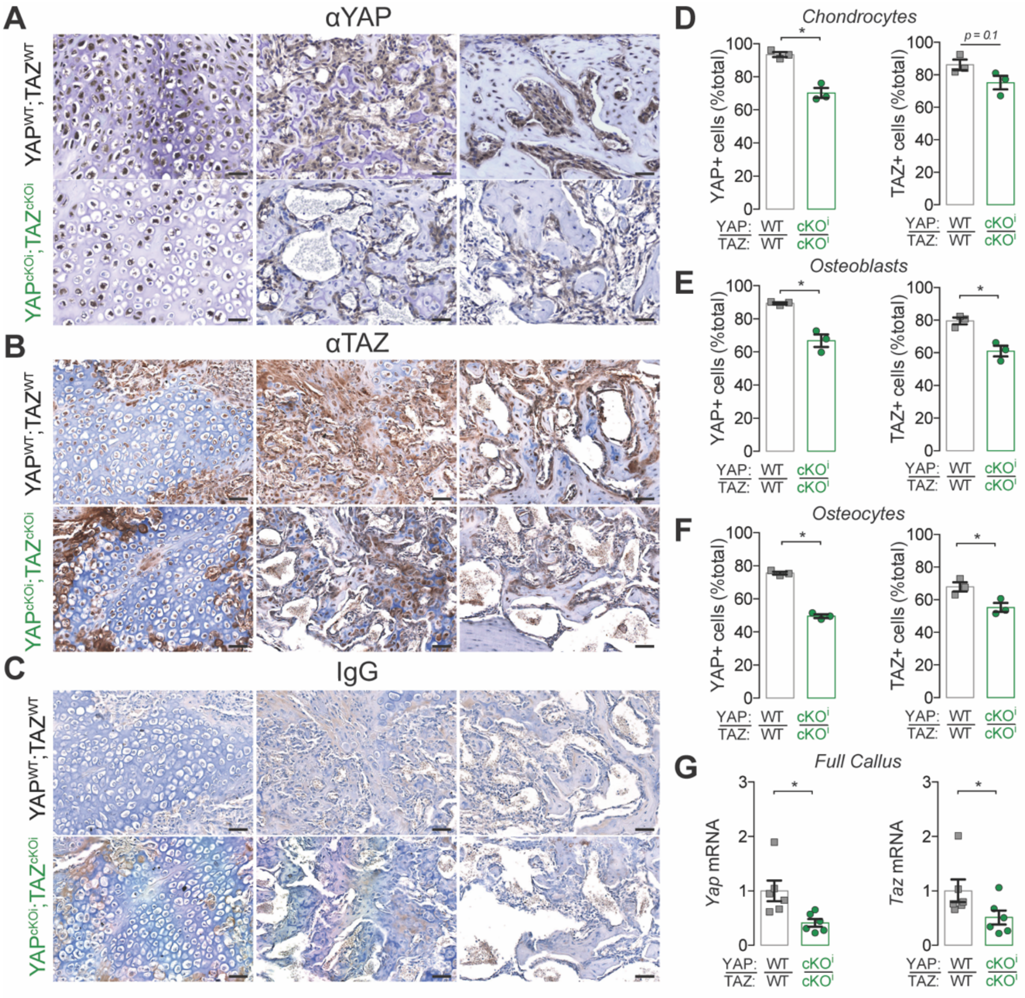


**Supplemental Fig. 3. Adult onset-inducible, homozygous YAP/TAZ deletion from Osterix-expressing cells moderately reduced callus YAP/TAZ expression on 14 dpf. A-C)** Representative micrographs of chondrocyte, osteoblast, and osteocyte immunostaining for YAP, TAZ, and IgG control sections at 14 dpf. Quantification of YAP and TAZ protein expression in **(D)** chondrocytes, **(E)** osteoblasts, and **(F)** osteocytes from callus sections at D14. **G)** *Yap* and *Taz* mRNA expression, relative to *18S rRNA*, from callus bone preparations at 14 dpf. Data are presented as individual samples in scatterplots and bars corresponding to the mean and standard error of the mean (SEM). N = 6 per group for qPCR and N = 3 per group for IHC. Scale bars equal 50 μm for all images.

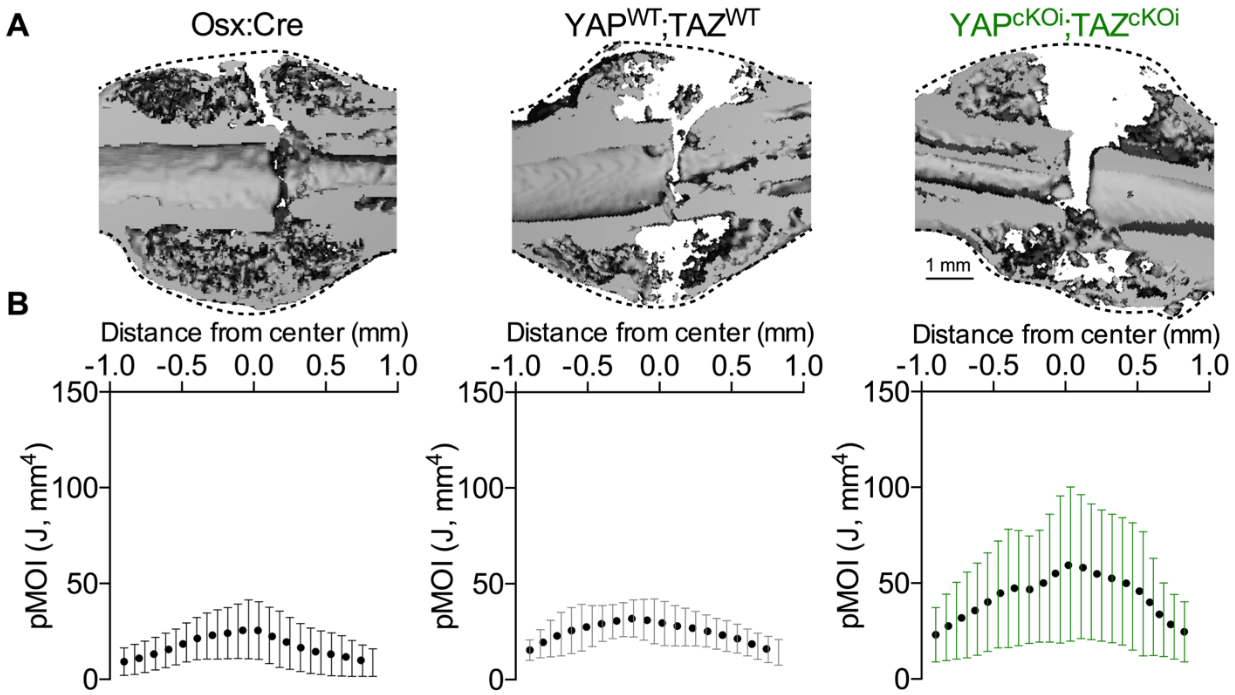


**Supplemental Fig. 4. Adult onset-inducible, homozygous YAP/TAZ deletion from Osterix-expressing cells increased variability in overall callus size. A)** 14 dpf microCT reconstruction of longitudinal cut planes within the callus. Dotted lines represent how all tissue within the callus were included in polar moment of inertia analysis **B)** Polar moment of inertia distributions of the entire callus for Osx:Cre, $YAP^{WT}$;$TAZ^{WT}$, and $YAP^{cKOi}$;$TAZ^{cKOi}$ genotypes. Data were binned into 25 equal distance bins from the center of the callus and presented as dots representing the mean and bars corresponding to the standard deviation (SD). Sample sizes, N = 5-8. Scale bars indicate 1 mm for microCT reconstructions.

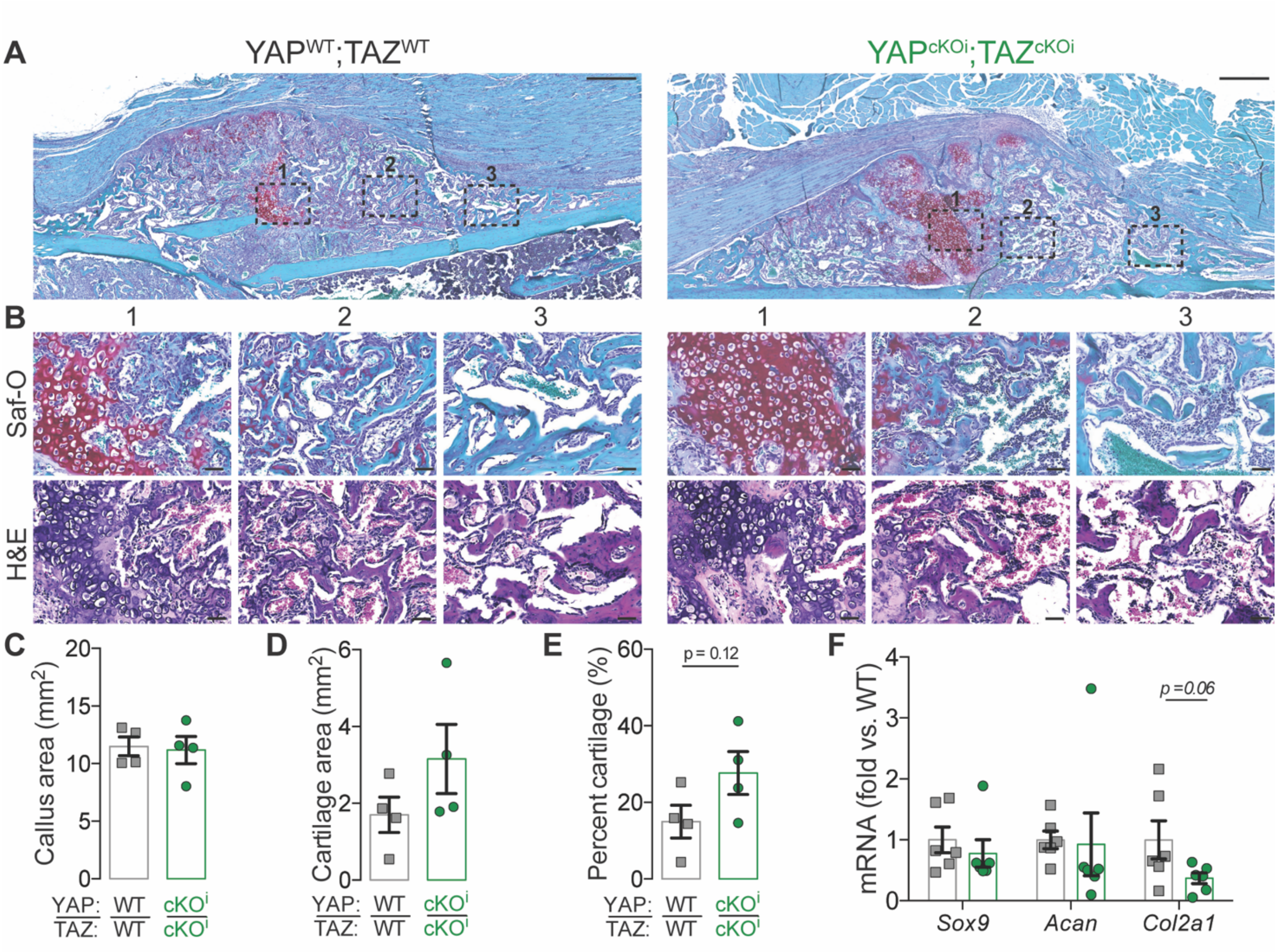


**Supplemental Fig. 5 Adult onset-inducible, homozygous YAP/TAZ deletion from Osterix-expressing cells did not significantly alter cartilaginous callus formation at 14 dpf. A)** Micrographs of Safranin-O stained calluses at 14 dpf. Black dotted boxes outline three zoomed-in regions of interest found for each genotype in **(B)**. Quantification of cartilaginous callus histomorphometry at 14 dpf of **(C)** total callus area, **(D)** cartilage area, and **(E)** percent cartilage area of total callus. **F)** *Sox9*, *Acan*, and *Col2a1* mRNA expression, relative to *18S rRNA*, from callus lysate preparations at 14 dpf. Data are presented as individual samples in scatterplots and bars corresponding to the mean and standard error of the mean (SEM). N = 6 per group for qPCR and N = 3-4 per group for histomorphometry. Scale bars equal 50 µm for all zoomed images and 500 µm for callus images.

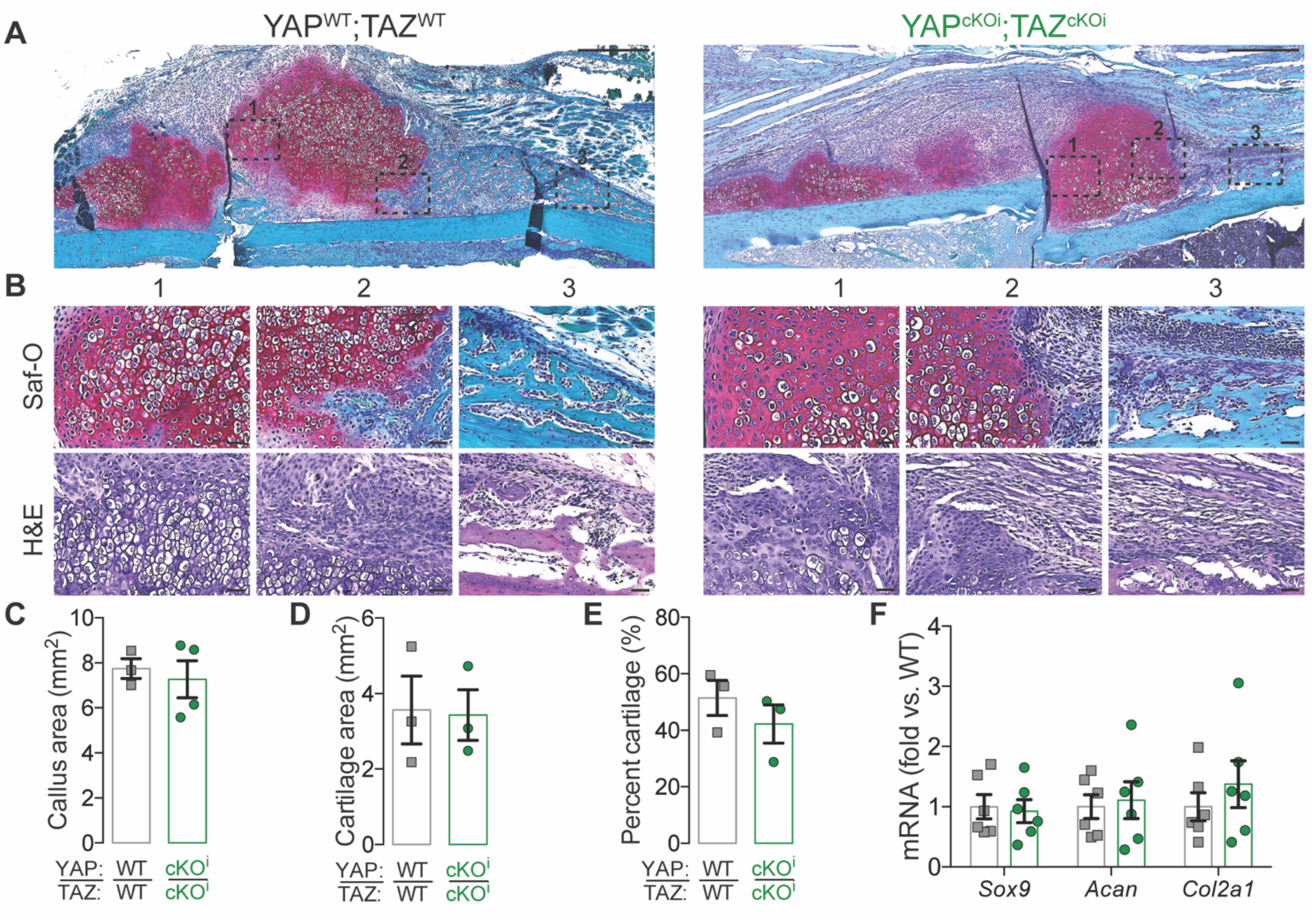


**Supplemental Fig. 6. Adult onset-inducible, homozygous YAP/TAZ deletion from Osterix-expressing cells did not significantly alter cartilaginous callus formation at 7 dpf. A)** Micrographs of Safranin-O stained calluses at 7 dpf. Black dotted boxes outline three zoomed-in regions of interest found for each genotype in **(B)**. Quantification of cartilaginous callus histomorphometry at 7 dpf of **(C)** total callus area, **(D)** cartilage area, and **(E)** percent cartilage area of total callus. **F)** *Sox9*, *Acan*, and *Col2a1* mRNA expression, relative to *18S rRNA*, from callus lysate preparations at 7 dpf. Data are presented with individual samples in scatterplots and bars corresponding to the mean and standard error of the mean (SEM). N = 6 per group for qPCR and N = 3 per group for histomorphometry. Scale bars equal 50 μm for all zoomed images and 500 μm for callus images.

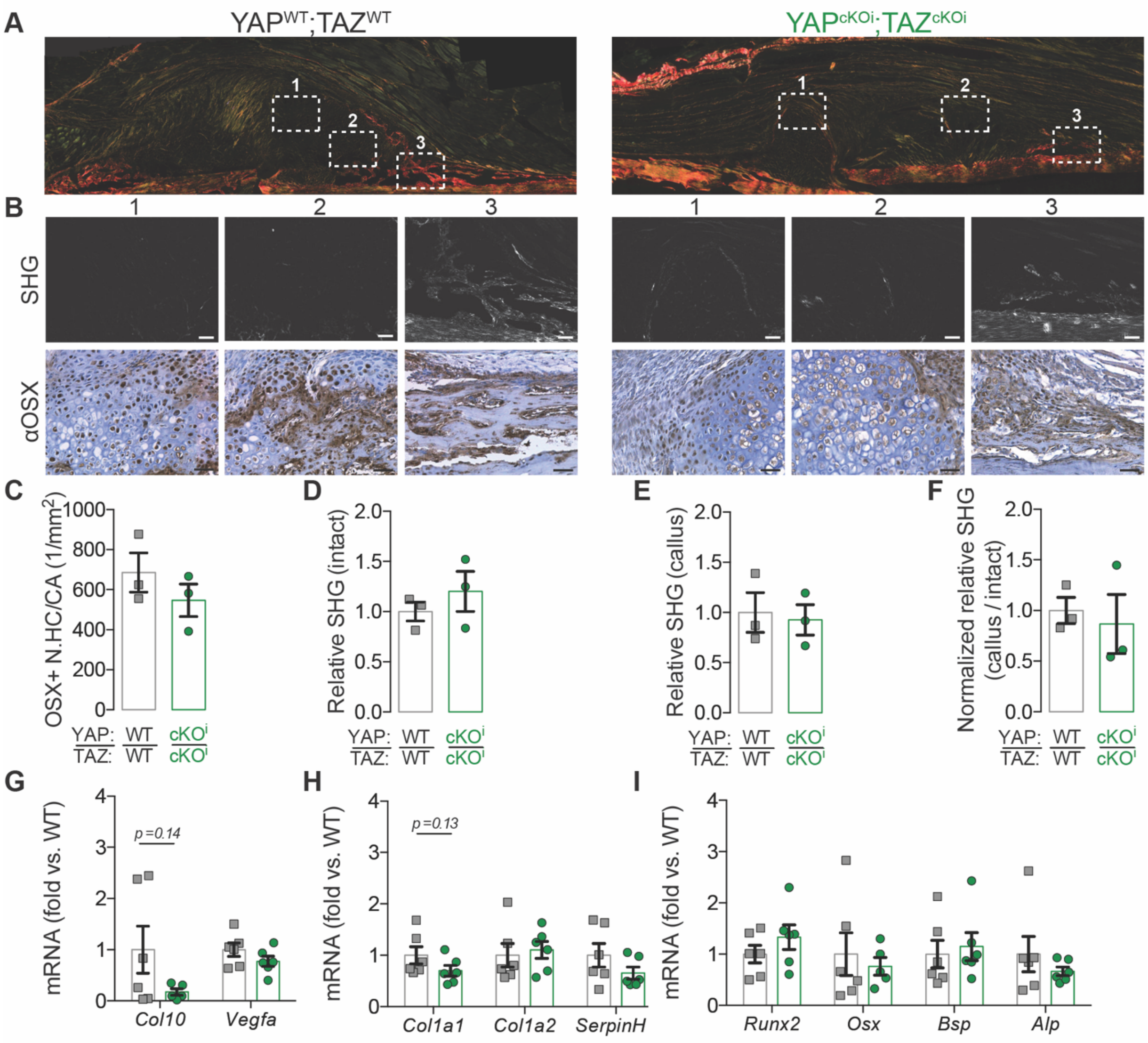


**Supplemental Fig. 7. Adult onset-inducible, homozygous YAP/TAZ deletion from Osterix-expressing cells did not significantly alter endochondral ossification or collagen matrix composition at 7 dpf. A)** Micrographs of Picrosirius Red stained calluses at 7 dpf. White boxes outline three zoomed-in regions of interest found for each genotype for both second harmonic generated imaging (SHG) and anti-Osterix (αOSX) immunostaining in **(B)**. Quantification of callus histomorphometry at 7 dpf of **(C)** the number Osterix-positive hypertrophic chondrocytes per cartilage area (OSX+ N.HC/CA). Quantification of relative SHG intensity per bone area at 7 dpf of **(D)** the intact cortical bone, **(E)** the newly formed callus bone, and **(F)** normalized callus-to-intact SHG intensity. **G)** *Col10* and *Vegfa* mRNA expression, relative to *18S rRNA*, from callus lysate preparations at 7 dpf. **H)** *Col1a1*, *Col1a2*, and *SerpinH1* mRNA expression, relative to *18S rRNA*, from callus lysate preparations at 7 dpf. **I)** *Runx2*, *Osx*, *Bsp*, and *Alp* mRNA expression, relative to *18S rRNA*, from callus lysate preparations at 7 dpf. Data are presented with individual samples in scatterplots and bars corresponding to the mean and standard error of the mean (SEM). N = 6 per group for qPCR and N = 3 per group for histomorphometry. Scale bars equal 50 μm for all zoomed images and 500 μm for callus images.

**Supplemental Table 1: qPCR primers.** Mouse primers used for qPCR

| Gene | | Primer Sequence (5' to 3') |
|---|---|---|
| *18S rRNA* | F<br>R | CGAACGTCTGCCCTATCAAC<br>GGCCTCGAAAGAGTCCTGTA |
| *Yap* | F<br>R | GATGTCTCAGGAATTGAGAAC<br>CTGTATCCATTTCATCCACAC |
| *Taz* | F<br>R | GGATACAGGTGAAAATTCCG<br>GATTACAGCCAGGTTAGAAAG |
| *Sox9* | F<br>R | AGTACCCGCATCTGCACAAC<br>ACGAAGGGTCTCTTCTCGCT |
| *Acan* | F<br>R | CCTGCTACTTCATCGACCCC<br>AGATGCTGTTGACTCGAACCT |
| *Col2a1* | F<br>R | GACTGAAGGGACACCGAG<br>CCAGGGATTCCATTAGAG |
| *Col10* | F<br>R | ATGCTGCCTCAAATACCCT<br>TGCCTTGTTCTCCTCTTACT |
| *Vegfa* | F<br>R | TAGAGTACATCTTCAAGCCG<br>TCTTTCTTTGGTCTGCATTC |
| *Runx2* | F<br>R | AGCCTCTTCAGCGCAGTGAC<br>CTGGTGCTCGGATCCCAA |
| *Col1a1* | F<br>R | GCTCCTCTTAGGGGCCACT<br>CCACGTCTCACCATTGGGG |
| *Col1a2* | F<br>R | GCTCCTCTTAGGGGCCACT<br>CCACGTCTCACCATTGGGG |
| *SerpinH1* | F<br>R | AGCCGAGGTGAAGAAACCC<br>CATCGCCTGATATAGGCTGAAG |
| *Osx* | F<br>R | CTGGGGAAAGGAGGCACAAAGAAG<br>GGGTTAAGGGAGCAAAGTCAGAT |
| *Alp* | F<br>R | GGACAGGACACACACACACA<br>CAAACAGGAGAGCCACTTCA |
| *Bsp* | F<br>R | ACAATCCGTGCCACTCACT<br>TTTCATCGAGAAAGCACAGG |
| *Cyr61* | F<br>R | CTGCGCTAAACAACTCAACGA<br>GCAGATCCCTTTCAGAGCGG |
| *Ctgf* | F<br>R | GGGCCTCTTCTGCGATTTC<br>ATCCAGGCAAGTGCATTGGTA |
